# Pervasive compartment-specific regulation of gene expression during homeostatic synaptic scaling

**DOI:** 10.1101/2020.08.31.274993

**Authors:** David Colameo, Marek Rajman, Michael Soutschek, Silvia Bicker, Lukas von Ziegler, Johannes Bohacek, Pierre-Luc Germain, Christoph Dieterich, Gerhard Schratt

**Affiliations:** Lab of Systems Neuroscience, Institute for Neuroscience, Department of Health Science and Technology, Swiss Federal Institute of Technology ETH, 8057 Zurich, Switzerland; Institute for Physiological Chemistry, Biochemical-Pharmacological Center Marburg, Philipps-University of Marburg, 35032 Marburg, Germany; Lab of Behavioural and Molecular Neuroscience, Institute for Neuroscience, Department of Health Science and Technology, Swiss Federal Institute of Technology ETH, 8057 Zurich, Switzerland; Institute for Neuroscience, Department of Health Science and Technology, Swiss Federal Institute of Technology ETH, 8057 Zurich, Switzerland; Lab of Statistical Bioinformatics, Department of Molecular Life Sciences, University of Zürich, 8057 Zurich, Switzerland; Section of Bioinformatics and Systems Cardiology, Department of Internal Medicine III and Klaus Tschira Institute for Integrative Computational Cardiology, University of Heidelberg, Germany; Neuroscience Center Zurich, ETH Zurich and University of Zurich, Switzerland

**Keywords:** homeostatic plasticity, synaptic scaling, local translation, cellular compartment, RNA-binding protein, microRNA

## Abstract

Synaptic scaling is a form of homeostatic plasticity which allows neurons to adjust their action potential firing rate in response to chronic alterations in neural activity. Synaptic scaling requires profound changes in gene expression, but the relative contribution of local and cell-wide mechanisms is controversial. Here we performed a comprehensive multi-omics characterization of the somatic and process compartments of primary rat hippocampal neurons during synaptic scaling. Thereby, we uncovered both highly compartment-specific and correlated changes in the neuronal transcriptome and proteome. Whereas downregulation of crucial regulators of neuronal excitability occurred primarily in the somatic compartment, structural components of excitatory postsynapses were mostly downregulated in processes. Motif analysis further suggests an important role for trans-acting post-transcriptional regulators, including RNA-binding proteins and microRNAs, in the local regulation of the corresponding mRNAs. Altogether, our study indicates that compartmentalized gene expression changes are widespread in synaptic scaling and might co-exist with neuron-wide mechanisms to allow synaptic computation and homeostasis.

## Introduction

Synaptic scaling is an important cellular mechanism that keeps excitatory synaptic strength in a physiological range in response to chronic (>6 hours) changes in synaptic input^1^. In the mammalian brain, synaptic scaling plays important roles during neural circuit development (e.g. activity-dependent development of the visual and barrel cortex)^2,3^ and in cognition in the adult (e.g. memory consolidation during sleep)^4,5^. Moreover, defects in synaptic scaling have been associated with the pathophysiology of several neurological diseases, e.g. autism (Rett Syndrome), mental retardation (e.g. Fragile-X Syndrome)^6^, epilepsy^7^ and mood disorders^8^. Recent evidence suggests that synaptic scaling requires changes in gene expression, both at the level of mRNA transcription^9,10^ and translation^11-14^. This leads to a remodelling of the synaptic proteome, in particular by synthesis/degradation of neurotransmitter receptors, presynaptic proteins and components of the calcium-dependent signalling pathways ^13-15^. The molecular mechanisms underlying these orchestrated changes in *de novo* protein synthesis, however, are only poorly understood. In hippocampal neurons, specific mRNAs are locally translated in the synapto-dendritic compartment, which provides neurons with a means to regulate the activity of individual synapses or at least dendritic branches^16^. Interestingly, in contrast to classical Hebbian synaptic plasticity, homeostatic plasticity was initially believed to operate exclusively at a global level, either at the level of entire neurons or networks^9^. However, recent theoretical considerations challenged this view and suggested an important contribution of local mechanisms (e.g. operating at the level of individual dendritic segments) to synaptic scaling^17^. This view is supported by experimental results from cultured hippocampal neurons, which show local dendritic regulation of retinoic acid receptor signalling^18^ and AMPA-type glutamate receptor synthesis^19,20^ in response to synaptic scaling up induced by chronic activity blockade. In contrast, whether synaptic downscaling in response to chronically elevated activity similarly involves local regulation of gene expression has not been addressed. However, compartmentalized changes in gene expression and their regulatory mechanisms during homeostatic plasticity or any other form of synaptic plasticity have not been comprehensively assessed on a more general level using multi-omics approaches.

## Results

In this study, we interrogated compartmentalized gene expression in rat hippocampal neurons undergoing homeostatic plasticity using a multi-omics approach. Therefore, we combined two *in vitro* model systems that we had previously established. First, a compartmentalized primary rat hippocampal neuron culture system, whereby neurons are plated on the upper side of filter inserts with small pore size (1 μm) that selectively allow the growth of neuronal processes (axons, dendrites), but not somata, to the lower side of the inserts (Fig. 1A)^21^. Second, a pharmacological stimulation protocol (48 hours treatment with the GABA-A receptor antagonist picrotoxin (PTX)) that induces a robust and chronic increase in network activity followed by downscaling of excitatory synaptic strength^22^. To validate the compartmentalized culture system, we first performed a comparative transcriptomic analysis of the somatic and process compartment in unstimulated neurons using ribosome depletion RNA sequencing (ribo(-)RNAseq). Thereby, we identified 1,231/1,673 genes that were significantly enriched with a logFC of greater than +/-1 in the somatic/process compartment, respectively (Fig. 1B). As expected, many well-known dendritically and axonally localized mRNAs, e.g. Camk2a, Shank2/3, Homer1, Dlg4 and Syn1, were significantly enriched in the process compartment. In contrast, genes encoding nuclear proteins, such as Polr2c, Sfpq and Nono, were significantly enriched in the somatic compartment (Fig. 1B). GO-term analysis indicates the overrepresentation of specific cellular components in these compartments: ribosomal subunits, postsynaptic density and mitochondrial respiratory chain complex in the process compartment; extracellular matrix, extracellular space and secretory granule lumen in the somatic compartment (Fig. 1C; Suppl. Fig. S1). Together, these results are consistent with previous large-scale studies^23-26^ and suggest that our approach is able to faithfully capture differences in mRNA distribution between neuronal compartments.

**Figure 1:**
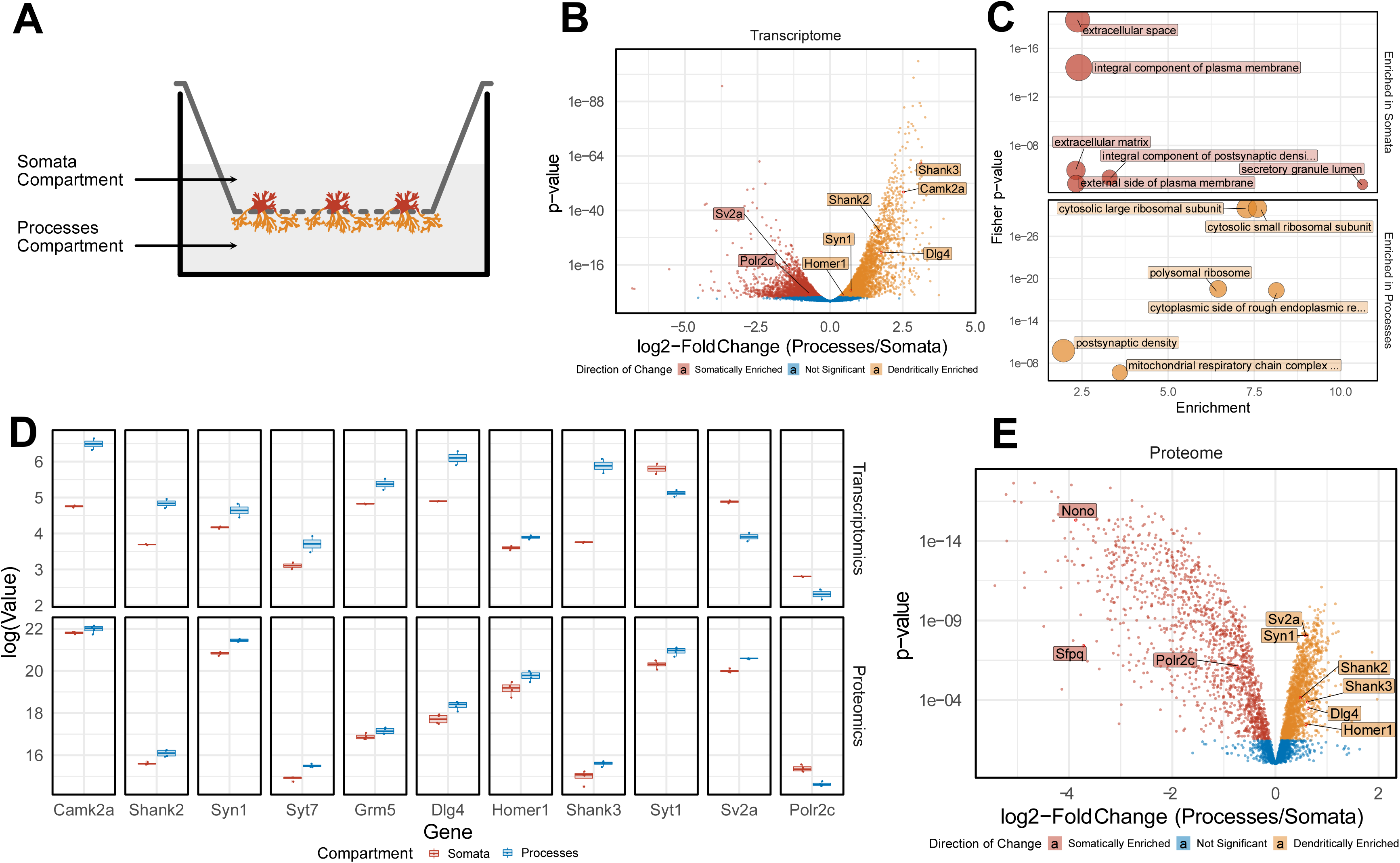
Compartment-specific localization of neuronal transcripts and proteins. (A) Schematic of the workflow for the culture of primary hippocampal neurons and 48 hours of PTX-treatment. (B) Volcano-plots demonstrating enrichment of transcripts in either the somatic (red) or process (yellow) compartment with representative genes highlighted. (C) Top Gene Ontology Pathway Analysis for transcripts enriched in the somatic and process compartment. (D) Boxplots of the highlighted genes enriched in either the somata or processes at the transcript or protein level. (E) Volcano-plots demonstrating enrichment of proteins in either the somatic (red) or process (yellow) compartment with representative genes highlighted.

We next investigated whether mRNA compartmentalization translated into corresponding changes in the neuronal proteome. Towards this aim, we prepared protein extracts from the somatic and process compartment of primary rat hippocampal neurons and subjected them to label-free proteomics (see material and methods). In four independent preparations, we identified unique peptides corresponding to a total of 4,250 different proteins in the somatic/process compartment, demonstrating a high sensitivity of our approach despite the low amount of starting material. Principal component analysis (PCA) of the replicates indicates a high reproducibility of both the RNA-seq and proteomics workflow (Suppl. Fig. S2). Similar to transcriptomics, a large number of neuronal proteins displayed preferential localization to either the somatic or process compartment (Fig. 1D). For the majority of differentially localized mRNAs, the respective proteins showed a corresponding compartment-specific expression (e.g. Syn1, Shank2/3, Dlg4, Homer1, Polr2c) (Fig. 1E). This suggests that local mRNA translation in processes contributes to the compartment-specific expression of those genes. However, for some genes (e.g. the presynaptic Sv2a and Syt1), mRNA and protein enrichment did not match, suggesting that protein transport plays a major role in the subcellular localization of these genes.

Having validated our cell culture model, we went on to investigate dynamic compartment-specific changes in the transcriptome/proteome upon synaptic downscaling in response to 48 h PTX application. Using Ribo(-)-RNAseq, we identified hundreds of RNAs that were differentially expressed (q<0.05) between PTX-and mock-treated conditions in the somatic (n=972; 431 down / 541 up) and process (n=949; 525 down / 424 up) compartment (Fig. 2A). Of those, about 50% (463) were common to both compartments, demonstrating that a large fraction of PTX-responsive RNAs displays compartment-specific regulation (Fig. 2B). To better visualize compartment-specific effects of PTX, we plotted logFC in somata vs. logFC in processes and colour-coded gene groups according to their regulation (Fig. 2C). Examples of candidate mRNAs within each of these gene groups are marked in Fig. 2C, e.g. process up (Sort1), process down (Srcin1, Add2, Dnajc6), as well as genes commonly up-(Plk2) or downregulated (Camk2a, Atp2b4, Shank2) in either compartment. In addition to protein-coding mRNAs, PTX also altered the expression of many non-coding RNAs in a compartment-specific manner, e.g. long non-coding RNAs (lncRNAs) and primary microRNAs (miRNAs) (suppl. Fig. S3). To obtain insight into the biological function of compartment-specific regulation by PTX, we performed GO term analysis. (Fig. 2D; suppl. Fig. S4). Genes specifically upregulated in the process compartment were often associated with the cell cycle (e.g. Cdc20) or metabolism, whereas “somata up genes” were enriched in neuropeptide signalling (e.g. Tac1) and intracellular trafficking (e.g. Sort1). Consistent with PTX-mediated downscaling of excitatory synapses, PTX-downregulated genes in both compartments were highly enriched for regulators of excitatory synaptic transmission. However, whereas the majority of “somata down genes” encode for regulators of synaptic excitation, e.g. ionotropic glutamate receptors (subunits of the AMPA-, NMDA- and Kainate-type) and transmembrane transporters (e.g. several members of the Slc family), “process down genes” mostly encode for structural components of the postsynaptic density, e.g. scaffolding (e.g. Shank1,3, Dlg3, Srcin1, Syngap1, Dlgap1, Homer2/3) and cytoskeletal proteins (e.g. Add2, Dnm1, Map1a/b/s). Taken together, our results demonstrate that compartment-specific regulation of the neuronal transcriptome during synaptic scaling is more pervasive than previously anticipated. We went on to validate selected high-ranking candidates that display robust compartment-specific regulation by PTX, particularly focusing on genes that are preferentially regulated in the process compartment. Using qPCR, we validated PTX-dependent differential regulation of several genes, such as Sort1, Add2, Shank2 and Srcin1 (Fig. 3A). Furthermore, we detected upregulation of Plk2 in either compartment, consistent with a previous report^27^. Compartment-specific regulation by PTX was further confirmed for Add2, Sort1 and Dnajc6 at the single neuronal level using single-molecule fluorescent *in situ* hybridization (smFISH) (Fig. 3B, C). smFISH further allowed us to distinguish between dendritic and axonal processes. Whereas PTX selectively reduced dendritic Add2 and Dnajc6 mRNA puncta, Sort1 mRNA puncta were significantly increased in dendrites. For all three genes, no significant changes were observed for somatic mRNA puncta upon PTX treatment.

**Figure 2:**
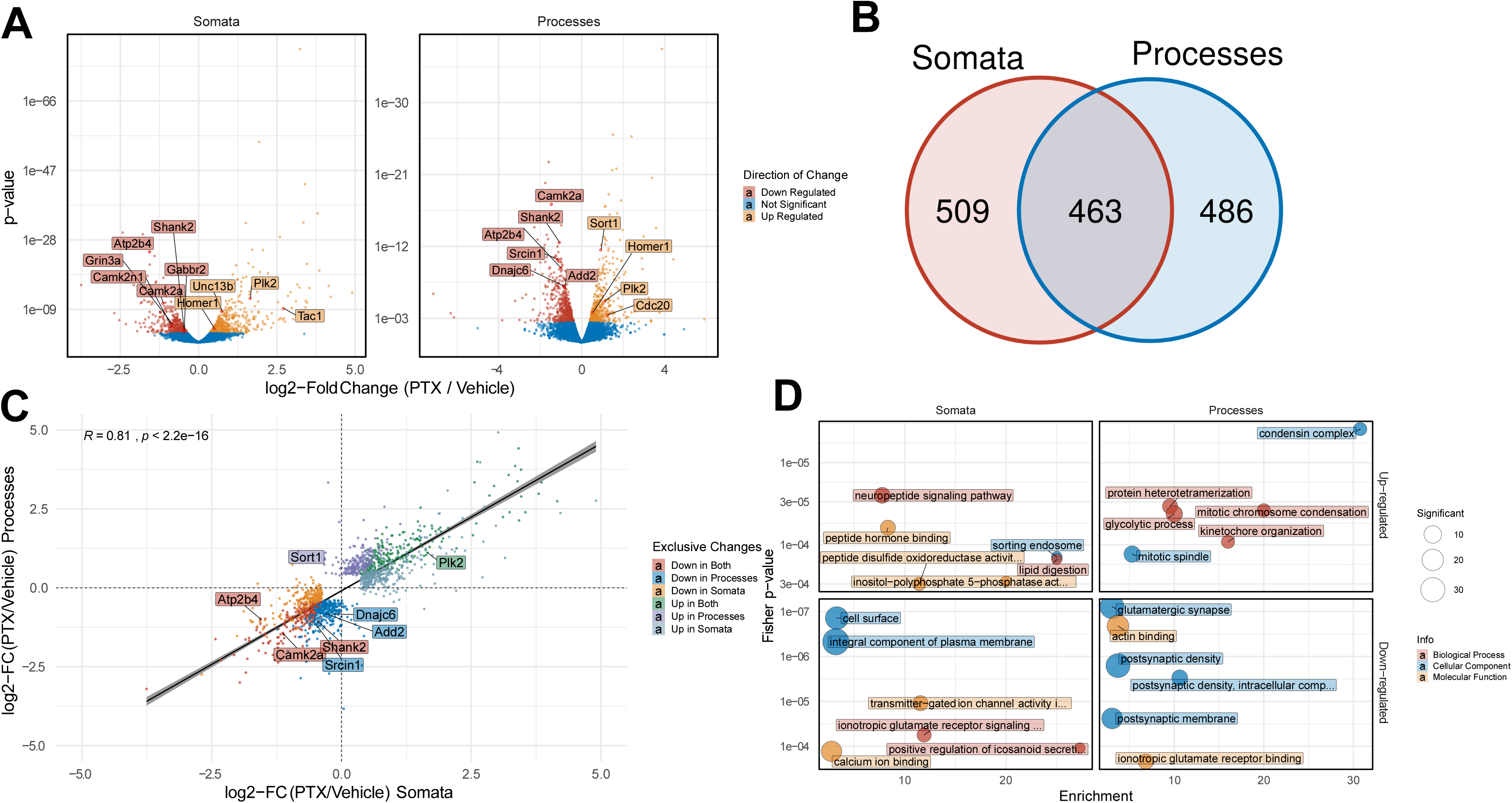
Compartment-specific regulation of neuronal transcripts by PTX. (A) Volcano plots representing transcript down-(red) or up-regulation (yellow) after 48 hours PTX in the somatic and process compartment. (B) Venn-Diagram of differentially expressed transcripts between somatic and process compartment (C) Pearson’s Correlation between log2-fold changes of differentially expressed genes significant in either the somatic or process compartment color-coded for exclusive changes in the respective compartments. (D) Top 6 Gene Ontology Pathways enriched in differentially expressed genes in either the somatic or process compartment color-coded for Cellular Component (blue), Molecular Function (yellow) or Biological Process (red).

**Figure 3:**
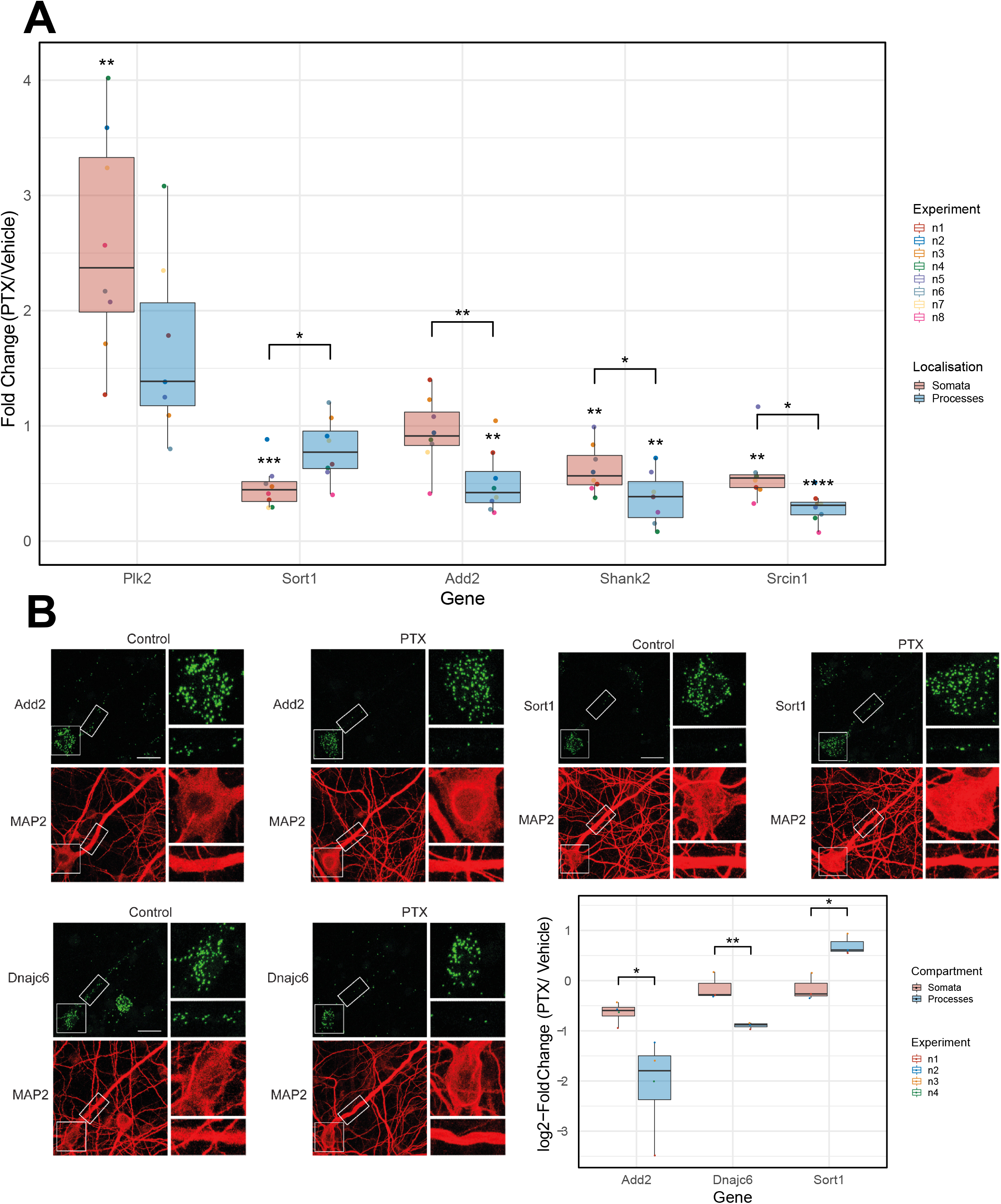
Validation of compartment-specific regulations. (A) Real-time quantitative PCR (RT-qPCR) of transcripts changing differentially between compartments using the compartmentalized cultures after 48 hours PTX-treatment. Boxplots represent median of 8 independent experiments (n=8); PTX-effect per compartment was assessed by one-sample Student’s t-test with μ0=1 multiple-test corrected using the Benjamin-Holm method; Differential compartment-effect was assessed by two-way ANOVA followed by Tukey’s post-hoc multiple comparison test; *p < 0.05; **p < 0.01; ***p < 0.001; ****p < 0.0001. (B) Representative images of single-molecule FISH (smFISH) in either control or 48 h PTX-treated rat hippocampal neurons (DIV20) using probes specific for Add2, Dnajc6 and Sort1 (green). MAP2 immunostaining (red) was used to visualize neuronal somata and dendrites. Inserts at higher magnification illustrate PTX-dependent changes in dendritic RNA puncta. Scale bar = 10 μm. (C) Quantification of B presented as boxplots. (n=3-4 with 8-10 cells averaged per condition and experiment; two-sample Student’s t-test; *p < 0.05; **p < 0.01).

We next wondered whether the PTX-induced compartment-specific changes in transcript levels translated into corresponding changes in protein. Although dynamic time- and polarity-dependent changes in the neuronal proteome have been previously demonstrated in synaptic scaling using labelled proteomics^12-14^, these studies did not address potential compartment-specific effects. Therefore, we extracted proteins from PTX-treated compartmentalized hippocampal neuron cultures and quantified proteome-wide changes using label-free proteomics. In total, we detected peptides corresponding to 4,250 different proteins with our approach. Differential expression analysis revealed that 660 proteins were significantly changing in somata and 320 proteins in processes, respectively (FDR<0.5, Fig. 4A; suppl. Fig. S5). Similar to our observations from RNA-seq, we detected substantial compartment-specific regulation of protein levels during synaptic downscaling, with only 162 proteins commonly regulated in both the somatic and process compartment (Fig. 4B). Similar to RNA-seq, GO-term analysis of differentially expressed proteins indicated a strong enrichment of synaptic proteins among PTX-downregulated proteins (suppl. Fig. S6). Importantly, although many of the top-ranking PTX-regulated mRNAs were not detectable by proteomics, PTX-dependent changes in RNA and protein levels in both compartments strongly correlated (Fig. 4C), e.g. preferential downregulation of Shank1, Dlg3, Crtc1 and Map1a in the process- and Grin2b, Slc2a13 in the somatic compartment, respectively. A correlation of PTX-dependent changes in RNA and protein levels is also observed for most of the candidates validated by qPCR (Fig. 4D). However, for a few genes, PTX-dependent changes in protein and RNA levels did not correlate. For example, Sort1 is significantly upregulated at the RNA level (Fig. 3C), but decreased at the protein level in the process compartment (Fig. 4D). This result suggests that additional regulatory mechanisms at the level of mRNA translation and/or protein turnover contribute to PTX-dependent process expression of those genes.

**Figure 4:**
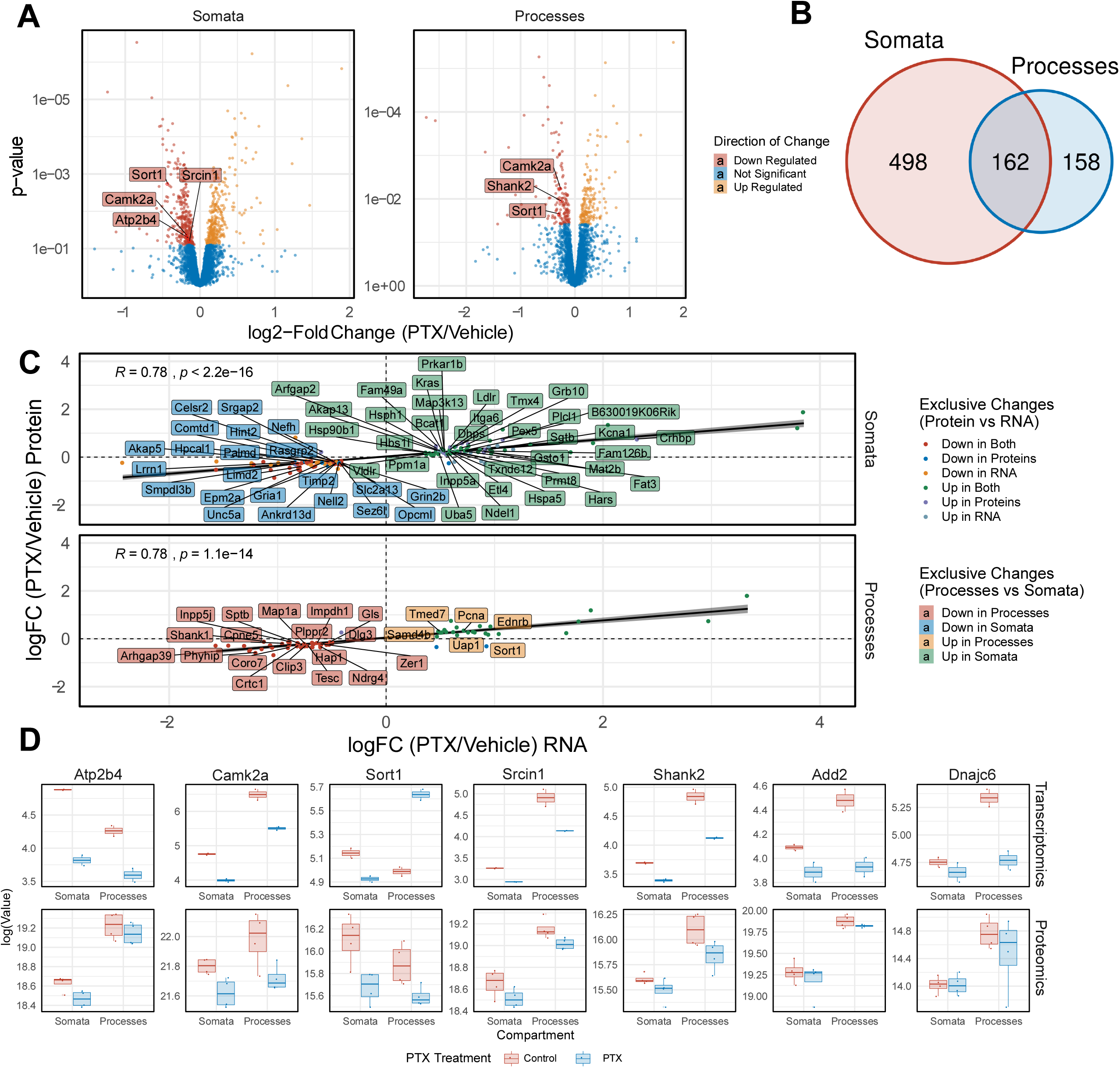
Compartment-specific regulation of neuronal proteins by PTX. (A) Volcano plots representing proteins down-(red) or up-regulation (yellow) after 48 hours PTX in the somatic and process compartment. FDR < 0.5 (B) Venn-Diagram of differentially expressed proteins between somatic and process compartment. (C) Pearson’s Correlation between log2-fold changes of differentially expressed genes changing at the transcript and protein-level. (D) Representative boxplots of genes changing after PTX at the transcript and protein level.

Next, we sought to characterize the molecular mechanisms which underlie the compartment-specific regulation of mRNA levels by PTX. Untranslated regions (UTRs) of mRNAs are considered as important determinants of both mRNA localization and post-transcriptional regulation. 3’UTRs in particular contain sequence elements for trans-acting post-transcriptional regulators, such as RNA-binding proteins (RBPs) and miRNAs (Fig. 5A). We first determined the average 3’UTR length in genes regulated in a compartment-specific manner (Fig. 5B). 3’UTRs of PTX-downregulated genes were slightly longer in comparison to upregulated genes in both the somatic and process compartment, although the difference was only statistically significant in the former compartment. Thus, 3’UTR length differences alone unlikely explain the high degree of compartment-specific gene regulation during synaptic scaling.

**Figure 5:**
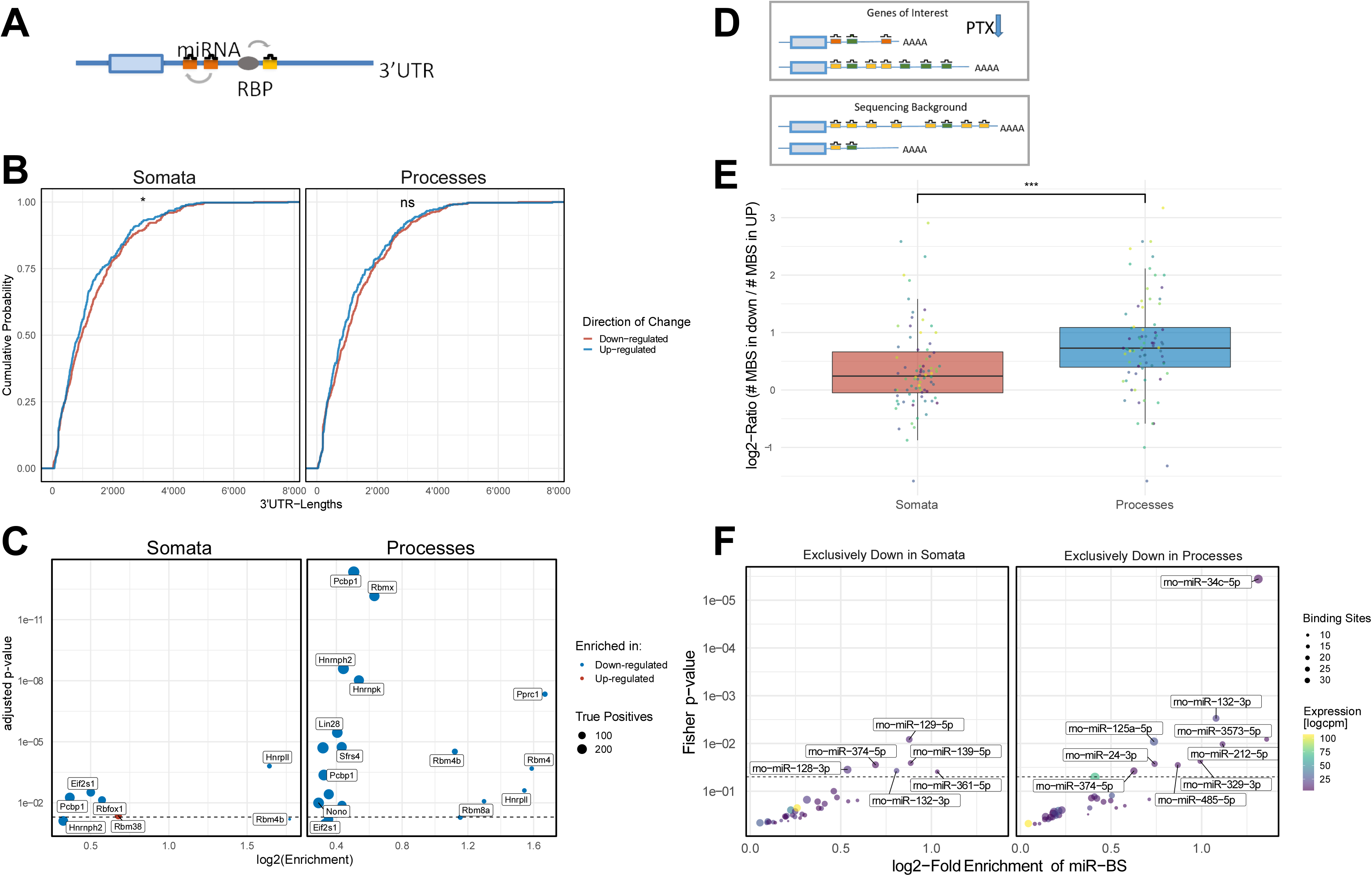
3’ UTR dependent mechanisms of regulation by RBPs and miRNAs: (A) Schematic of trans-acting RNA-binding proteins (RBPs) and microRNAs (miRNAs) (to be finalized) (B) Cumulative distribution of 3’-UTR lengths of down- and up-regulated genes in either compartment. Two-sample Kolmogorov-Smirnov test; ns > 0.05, *p-value < 0.05. (C) Enrichment analysis for RBP motifs in down- and up-regulated genes in either compartment. Fisher’s exact test was used to perform enrichment analysis; FDR-corrected. (D) Schematic of miRNA-binding site (MBS) enrichment (to be finalized). (E) Log2-Ratios between number of miRNA-binding sites found in the pool of up-versus down-regulated genes. Non-parametric Mann-Whitney U-test was used to test difference (mean somata: 0.37 vs mean processes: 0.783); ***p-value < 0.0001; n=80 miRNAs. (F) Enrichment of MBS in genes that are exclusively down-regulated in somata or processes compared to the sequencing background. The significant miRNA families (p-value < 0.05 and log2-fold enrichment > 0.5) are labelled by a representative miRNA. See Methods for detailed statistical analysis.

Subsequently, we investigated the compartment-specific enrichment of specific sequence motifs located within 3’UTRs. We first interrogated overrepresentation of RBP binding motifs determined in a previous large-scale *in vitro* study^28^ in the 3’UTRs of PTX-regulated mRNAs (Fig. 5C). Whereas only 5 motifs were significantly (Adjusted p-value <0.05) overrepresented in PTX-regulated somatic mRNAs, 16 motifs fulfilled these criteria in PTX-regulated process-enriched mRNAs (Fig. 5C). Interestingly, only one motif in total could be recovered from PTX-upregulated genes, suggesting that RBPs acting by 3’UTR-dependent mechanisms are mostly involved in PTX-mediated gene repression. Among the motifs most highly enriched in PTX-downregulated genes in somata is the Rbfox1 consensus binding site (GCAUG). This fits well with our previous results showing an important role of Rbfox1 in synaptic scaling at the level of the entire neuron ^15^. Concerning process-enriched RBP motifs, many of them are recognized by known regulators of mRNA stability and translation, such as several members of the hnRNP family of RBPs (e.g. hnRNP-K^29-31^, Nono^31^ and Lin-28^32^). This suggests an important contribution of RBPs to compartment-specific gene regulation during synaptic scaling.

miRNAs represent an additional class of gene-specific regulatory molecules that negatively affect mRNA expression by recruiting a silencing protein complex to specific sites (miRNA binding sites, MBS) in 3’UTRs^33^. To estimate miRNA contribution to PTX-dependent regulation, we performed an MBS enrichment analysis. Briefly, the number of MBS within a gene set of interest (i.e. PTX-downregulated genes in the process compartment) were compared to those present in the sequencing background (Fig. 5D; see Methods for further details). We calculated enrichment scores for MBS of neuronally expressed miRNAs (n=221) in the 3’UTR of genes that are downregulated by PTX in a compartment-specific manner. In both compartments, MBS were on average more abundant in PTX down-compared to upregulated genes (log2(PTX/control) > 0; Fig. 5E), as expected. However, the enrichment value (log2-fold ratio) obtained from processes was significantly higher compared to somata (Fig. 5E; p<0.001). Taken together, these results suggest that miRNA-dependent mRNA repression plays a particularly important role in the process compartment during synaptic downscaling. At the level of individual miRNA families, specific sets of MBS were significantly enriched over background in PTX-downregulated genes in either the somatic or process compartment (Fig. 5F). Our approach independently identified miR-129-5p, which we previously showed to be required for synaptic downscaling^15^, as the miRNA whose binding motifs were most strongly enriched in PTX-downregulated genes in the somatic compartment. This finding indicates that our approach is able to identify functionally important miRNAs solely on the basis of differential expression analysis. On the other hand, our enrichment analysis identified an overrepresentation of several miRNA binding motifs in mRNAs that are specifically downregulated by PTX in the process compartment, such as the activity-regulated miR-132/212^34^, miR-24-3p, miR-125a-5p, as well as the miR-379-410 cluster members miR-329-3p and miR-485-5p^35^. One miRNA that appears to be specifically active in dendrites during PTX-induced downscaling is miR-34c-5p, a member of the miR-34/449 family (Fig. 5F, suppl. Fig. S7). A total of 53 PTX-downregulated genes in processes contain miR-34/449 binding motifs in their 3’UTRs, including important synaptic regulatory proteins such as Add2, Shank3, Cntn2, Cacng2, Scn2b, Mpp2 and Crtc1 (Torc1) (Fig. 6).

**Figure 6:**
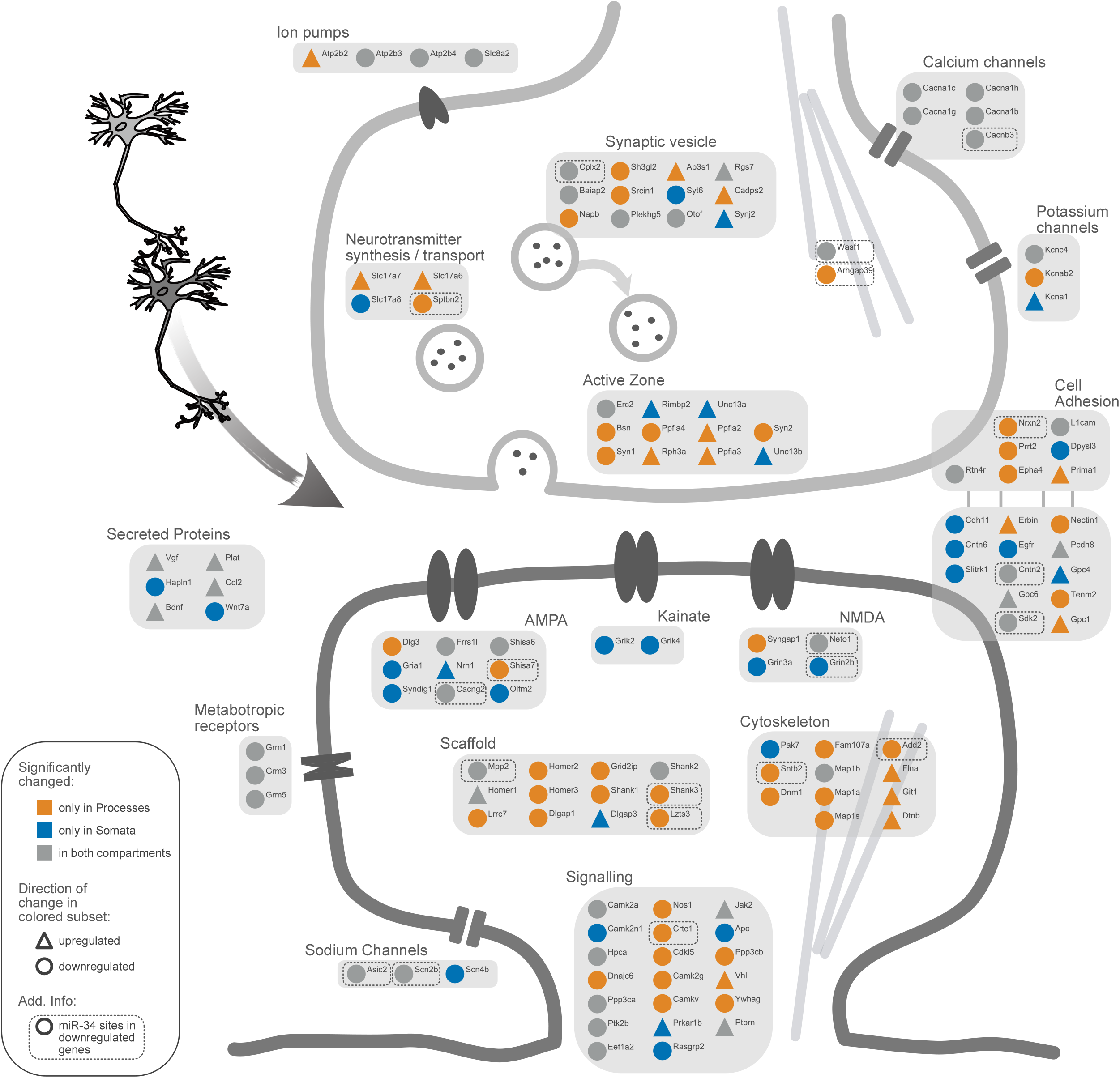
PTX-mediated changes in synaptic genes. Schematic representation of significantly changing genes (compartment RNA-sequencing) associated with excitatory synapses. Genes differentially expressed in only one compartment are highlighted with either yellow (processes compartment) or blue (somata compartment). The direction of change is illustrated as symbol shape (circle = sign. downregulated genes, triangle = sign. upregulated genes). miR-34 family binding sites in sign. downregulated genes are marked with a dashed line.

Taken together, compartment-specific regulation of neuronal transcripts during synaptic downscaling is associated with specific 3’UTR features, including unique miRNA and RBP motifs. Post-transcriptional regulation by miRNAs and RBPs appears to be particularly widespread during PTX-mediated downregulation of synaptic genes in the process compartment.

## Discussion

In this study, we provided the first comprehensive analysis of compartmentalized gene expression during synaptic plasticity, more specifically in response to homeostatic synaptic downscaling induced by prolonged treatment of primary rat hippocampal neurons with the GABA-A-R blocker PTX. A major outcome of our study was the observation that PTX-treatment induces widespread compartment-specific changes in the neuronal transcriptome and proteome. The existence of local and global mechanisms in homeostatic plasticity has been known for some time^36^, but a clear picture of how homeostatic feedback is structured at the molecular level has not yet emerged. Our study lends further support to the hypothesis that local synaptic and neuron-wide mechanisms co-exist in neurons and act in concert to counteract excessive excitation which would inevitably occur if feedforward Hebbian plasticity operated unconstrained^17^.

What distinguishes the somatic and process compartment during synaptic scaling? Some clues might come from our GO-term analysis, which showed a strong overrepresentation of genes involved in the regulation of neuronal excitability (e.g. ion channels, transporters, receptors) in the cell body, whereas the majority of regulation of structural synaptic genes (e.g. postsynaptic scaffolds, cytoskeletal proteins) occurred in the process compartment (Fig. 6). Consistent with this observation, many of the process-regulated mRNAs are known dendritic mRNAs, e.g. Shank1/3 and Homer2 and Syngap1, and have recently been shown to be preferentially translated in the synaptic neuropil *in vivo*^37^. On the other hand, “neuronal excitability genes” encode mostly transmembrane proteins, for which the somatic biosynthesis machinery might represent the major source, although dendritic translation has been demonstrated for some of these genes. Thus, mRNA sorting under basal conditions might already pre-determine compartment-specific responsiveness during scaling or other plasticity-inducing cues.

What are the molecular mechanisms underlying the compartment-specific regulation during synaptic scaling? In principle, compartment-specific mRNA changes could be a result of altered mRNA transcription, transport, stability and translation, or combinations thereof. However, several lines of evidence suggest that increased mRNA degradation might be the major driver for the observed downregulation of synaptic gene expression. First, mRNA and protein levels of differentially expressed genes strongly correlated, arguing against a major contribution of mRNA translation (Fig. 4C). Second, we did not obtain evidence for a PTX-dependent re-location of candidate mRNAs (e.g. Add2) between compartments in smFISH (Fig. 3B), making PTX-regulated RNA transport unlikely. In the future, assays that allow a direct visualization of local mRNA translation (e.g. Puro-PLA^38^) or local mRNA degradation (e.g. TREAT^39^) will have to be performed.

The striking overrepresentation of motifs for miRNAs and RBPs in 3’UTRs of PTX-regulated genes which are selectively downregulated in processes (Fig. 5C, F) further suggests that gene-specific post-transcriptional mechanisms might play a particularly important role in the local control of synaptic gene expression. With regard to miRNAs, many of the PTX-downregulated synaptic genes (e.g. Add2, Shank3, Cntn2, Cacng2, Scn2b, Mpp2) contain binding sites for the miR-34/449 family. A synaptic function of miR-34 was very recently reported in *Drosophila*, where it controls pre- and postsynaptic function at the neuromuscular junction (NMJ)^40^. miR-34 has also been implicated in stress-associated disorders, e.g. anxiety and epilepsy^41^, providing potential links between miR-34, synaptic scaling and neurological disease which should be explored in the future. Concerning RBPs, motifs for several members of the hnRNP family, Lin-28 and Nono were specifically enriched in “process down genes”, consistent with their reported presence in dendritic RNA granules^42^. Among them, hnRNP-K might be a strong candidate for follow-up experiments. Since hnRNP-K has mostly stabilizing functions on mRNA^43-45^, one possible scenario is that it counteracts miRNA activity under basal conditions and is subsequently inactivated upon scaling. In a broader context, specific combinations of motifs for trans-acting regulatory factors (“regulons”) might underlie the spatiotemporal control of neuronal gene expression in response to extracellular cues.

What could be the biological relevance of compartmentalized gene expression during homeostatic scaling? One can speculate that it might allow neurons to maintain excitability while taking into account individual plasticity needs of synapses or dendritic domains^46^. Intriguingly, a very recent study described the local allocation of proteins in so-called “synaptic neighborhoods”, about 10 micron long dendritic domains, in a related form of homeostatic plasticity, synaptic upscaling^47^. In agreement with our results, many of these allocated proteins might arise from local dendritic mRNA translation. Thus, local or quasi-local mechanisms operating at the level of synapses, neighbourhoods or dendritic domains might represent a common feature of homeostatic plasticity in excitatory neurons.

In conclusion, system-wide approaches like the one presented here might help to disentangle the spatiotemporal logic of the complex regulatory networks underlying synaptic plasticity and homeostasis and their aberrations in diseases such as epilepsy and autism.

## Methods

### Cell culture, transfection and stimulation

Primary cultures of Sprague–Dawley rats (Charles River Laboratories, Sulzfeld, Germany) embryonic hippocampal neurons were prepared as described previously^48^ and plated onto porous membrane cell culture inserts as described previously^21^. For stimulation, 18DIV neurons were treated either with Picrotoxin (PTX; 100 μM final concentration, Sigma) or vehicle (ethanol absolute) for 48h and lysed at 20DIV.

### Single molecule fluorescence *in situ* hybridization (smFISH)

Dissociated hippocampal neurons were fixed at 20DIV using 4% paraformaldehyde/4% sucrose/PBS for 30min at room temperature. FISH was performed using the QuantiGene (QG) ViewRNA kit (Affymetrix) according to the manufacturer’s protocol using probes for Add2, Dnajc6 and Sort1. The protease treatment step was omitted in order to maintain dendritic integrity. After completion of the FISH protocol, cells were processed for immunostaining, using an anti-MAP2 antibody (Sigma M9942; 1:1000) in GDB buffer (0.02% gelatin–0.5% Triton X-100– PBS). Images were acquired on a Leica SP5 laser-scanning confocal microscope. For z-stack images of whole cells, 12 consecutive optical sections were taken at a 0.4 μm interval with a resolution of 1024×1024 pixels using a 63x objective and a digital zoom factor of 2.5. For images of cell bodies, 15 consecutive optical sections were taken at a 0.4 μm interval with a resolution of 512×512 pixels using a 63x objective and a digital zoom factor of 5. For all pictures, the pinhole was set to 1 AU. Laser settings were kept constant between experimental conditions. Maximum intensity projections of the z-stacks were used for signal quantification, which was conducted in a blinded manner. The density of RNA particles was analysed by CellProfiler^49^, using the MAP2 immunostaining as mask to define cell bodies and dendrites.

### RNA extraction

RNA was extracted from primary hippocampal neuronal cultures using either peqGOLD Isolation Systems TriFast™ (Peqlab) to prepare sequencing library or mirVana™ total RNA Isolation Kit for quantitative real-time PCR following the manufacturer’s protocol. To remove potential DNA contamination, RNA samples were treated with TURBO™ DNase (Ambion) and RNA was re-extracted as described before. Samples were stored at −80°C until further use. Prior to RNA extraction, compartmentalized cell culture system was pre-processed as described^50^.

### Quantitative real-time PCR

RNA was reverse-transcribed with iScript™ reverse transcription supermix (Bio-Rad) using random hexamers (total RNA) or oligo dT20primers (poly-RNA) according to manufacturer’s instructions. Quantitative real-time PCR was performed with the Step One Plus Real-Time PCR System (Applied Biosystems), using iTaq SYBR Green Supermix with ROX(Bio-Rad) for detection of mRNA.

### Ribo(-) RNA-sequencing

Library preparation and sequencing were performed by the Max Planck-Genome-Centre Cologne, Germany (http://mpgc.mpipz.mpg.de/home/) for compartmentalized RNA samples. For the first step, rRNA depletion (Ribo-Zero rRNA removal Kit (Illumina)) has been performed with 1 μg total RNA per sample (n=2 samples per condition), followed by library preparation with NEBNext Ultra™ Directional RNA Library Prep Kit for Illumina (New England Biolabs). Sequencing was performed as 100 bp single read sequencing on HiSeq2500™ (Illumina).

### Protein Extraction

Primary hippocampal compartmentalized cultures were prepared as described earlier and treated at DIV19 with either 100μM PTX or vehicle for 48 hours. Inserts were then rinsed twice with ice-cold phosphate buffered saline (PBS, Gibco™) and lysed using a cell scraper in home-made RIPA-buffer (150mM NaCl, 1% Triton X-100; 0.5% Sodium Deoxycholate; 1mM EDTA; 1mM EGTA; 0.05% SDS; 50mM Tris pH8) containing a protease inhibitor cocktail (1:1000; Roche). Cell lysates were then homogenized by pipetting and centrifuged for 30min at 13’000xg (4°C). The supernatant was snap-frozen in liquid nitrogen and stored at −80°C until further use. Pierce™ BCA Protein Assay Kit (Thermo Scientific™) was used to quantify protein concentrations following the manufacturer’s protocol.

### Label-Free Proteomics

#### Protein digest and clean-up

Protein extracts were further processed with a filter assisted sample preparation protocol^51^. 20ug of protein were added to 30ul SDS denaturation buffer (4% SDS (w/v), 100mM Tris/HCL pH 8.2, 0.1M DTT). For denaturation, samples were incubated at 95°C for 5 min. Samples were diluted with 200ul UA buffer (8M urea, 100mM Tris/HCl pH 8.2) and then loaded to regenerated cellulose centrifugal filter units (Microcon 30, Merck Millipore, Billercia MA, USA). Samples were spun at 14000g at 35°C for 20 min. Filter units were washed once with 200ul of UA buffer followed by centrifugation at 14000g at 35°C for 15 min. Cysteines were alkylated with 100ul freshly prepared IAA solution (0.05M iodoacetamide in UA buffer) for 1 min at room temperature in a thermomixer at 600rpm followed by centrifugation at 14000g at 35°C for 10 min. Filter units were washed 3 times with 100ul of UA buffer then twice with a 0.5M NaCl solution in water (each washing was followed by centrifugation at 35°C and 14000g for 10 min). Proteins were digested overnight at room temperature with a 1:50 ratio of sequencing grade modified trypsin (0.4ug, V511A, Promega, Fitchburg WI) in 130ul TEAB (0.05M Triethylammoniumbicarbonate in water). After protein digestion over night at room temperature, peptide solutions were spun down at 14000g at 35°C for 15 min and acidified with 3ul of 20% TFA (trifluoroacetic acid).

#### Peptides Clean-up

Peptides were cleaned-up using StageTip C18 silica columns (SP301, Thermo Scientific, Waltham MA). Columns were conditioned with 200ul methanol followed by 200 ul of 60% ACN (acetonitrile) / 0.1% TFA. Columns were equilibrated with 2 x 150 ul of 3% ACN / 0,1% TFA. Samples were loaded onto the columns. They were then washed with 2 x 150 ul 3% ACN / 0.1% TFA and eluted with 150 ul 60% ACN / 0.1% TFA. Samples were lyophilized in a speedvac then re-solubilized in 19ul 3% ACN / 0.1% FA (formic acid) prior to LC-MS/MS measurement. 1ul of synthetic peptides (Biognosys AG, Switzerland) were added to each sample for retention time calibration.

#### LC-MS/MS measurements

Samples were measured on a QExactive (Thermo Fisher Scientific, Waltham MA, USA). Peptides were separated with an Eksigent NanoLC (AB Sciex, Washington, USA). We used a single-pump trapping 75-um scale configuration (Waters). 1ul of each were injected. Trapping was performed on a nanoEase™ symmetry C18 column (pore size 100Å, particle size 5um, inner diameter 180um, length 20mm). For separation, a nanoEase™ HSS C18 T3 column was used (pore size 100Å, particle size 1.8um, inner diameter 75um, length 250mm, heated to 50°C). Peptides were separated using a 120 min long linear solvent gradient of 5-35% ACN / 0.1% FA (using a flowrate of 300nl / min). Electronspray ionization with 2.6kV was used and a DIA method with a MS1 in each cycle followed by 35 fixed 20 Da precursor isolation windows within a precursor range of 400-1100 m/z was applied. For MS1 we used a maximum injection time of 200ms and an AGC target of 3e6 with a resolution of 60k in the range of 350-1500 m/z. MS2 spectra were acquired using a maximum injection time of 55ms an AGC target of 1e6 with a 30k resolution. A collision energy of 28 was used for fragmentation.

#### Protein search and quantification

We used Spectronaut™ (Biognosys, version 10) with directDIA for peak picking and sequence assignment. We used a *M. musculus* reference proteome for *R. norvegicus* from uniprot (UP000002494). We included a maximum of 2 missed cleavages, using a Tryptic in-silico digest with a KR/P cutting profile. Sequences in a range of 7-52 AA were considered. We included carbamidomethyl as fixed modification for cysteine, oxidation as variable modification for methionine and protein N-terminal acetylation as variable modification. Decoys were generated using a scrambled label free decoy method. A normal distribution estimator was used with a 1% FDR for q-value filtering. A maximum of 5 variable modifications were considered. Single hit was determined on the stripped sequence level. Major grouping was done by protein group ID and minor grouping by stripped sequence. Only proteotypic peptide sequences were considered and single hit proteins excluded. For the minor and major group quantification, the top 3 entries were used using the mean precursor/peptide quantity. A localized normalization strategy and interference correction were used. Machine learning was performed on a per run basis and iRT profiling was enabled.

### Bioinformatic analysis

#### RNA Sequencing analysis

Adapter sequences were trimmed from the reads using Trimmomatic 0.38 with ILLUMINACLIP 1:30:10, LEADING:5, TRAILING:5, SLIDINGWINDOW:5:15 AVGQUAL:20 and MINLEN:30. The reads were then mapped to the rat genome Rnor 6.0 using Subread subjunc 1.33.16 and the Ensembl 96 transcript annotation, and quantified with featureCounts (v.1.6)^52^. Counts were then aggregated to gene symbols and only genes with more than 20 reads in at least 2 samples for further processing. Counts were then aggregated to gene symbols and only genes with more than 20 reads in at least 2 samples were considered for further processing.

Differential expression analysis was performed with edgeR (v. 3.28)^53^ using TMM normalization. Specifically, linear models were fitted using glmFit through all 4 conditions (∼ *compartment* * *treatment*), and differentially expressed genes were identified by testing for the *treatment* effect in each compartment using separate contrasts.

For the enrichment analyses, a compartment specific background of expressed genes was determined by taking only genes with more than 20 reads in at least two conditions within the respective compartment into account.

#### Proteomics Data Analysis

A comprehensive library of detected proteins was generated by combining a search against a *R. Norvegicus* reference proteome with exclusively reviewed entries (Swiss-Prot, UP000002494, 8104 proteins) with a search against a reference proteome containing also predicted entries (TrEMBL, UniProt UP000002494, 21838 proteins) if proteins were undetectable in more than two samples. We detected 4047 proteins with no missing values across all conditions out of the total 4250 proteins. Values were first normalized by variance stabilizing transformation using the vsn package (version 3.54.0) and missing values were imputed using the MinProb function (q-value < 0.01) of the DEP package (version 1.8.0).

The package limma (version 3.42.2) was used to perform differential expression analysis using a linear multiplicative model fitted through all 4 conditions (∼ *compartment* * *treatment*) and differential expression was computed using empirical Bayes statistics to contrast *treatment*- and *compartment* effects.

#### GO enrichment analysis

Gene Ontology enrichment analysis was performed using the TopGo algorithm (v.2.40.0)^54^, essentially as described previously in^35^.

For the Compartment Analysis (Fig. 1, Suppl. Fig. 1), significantly enriched genes with a logFC of greater than +/-1 were tested against the respective specific background. Subsequently, the Top 6 / 10 GO-Terms in each ontology were plotted (with those filtered out that have more than 1000 annotated genes in CC or more than 300 annotated genes in BP).

PTX GO-terms were obtained by either testing exclusive gene sets (as characterized in Fig. 2c) against the respective specific background (Fig 2D) or by testing all significantly down-/ upregulated genes in a specific compartment (Suppl. Fig. 4). For Fig. 2D, the Top 5 obtained GO-Terms of all three ontologies were sorted by *Enrichment* and of those, the Top 6 GO-Terms plotted. GO-Terms with more than 1000 annotated genes in an ontology were filtered out.

GO-Terms enriched in the proteomics dataset were identified by aggregating multiple detected protein isoforms and subsequently performing enrichment analysis of significantly changing proteins (criteria proteomics data analysis) with TopGo against the background of detected proteins. For plotting, GO-terms with more than 500 annotated genes (CC) were filtered out.

#### miRNA binding site enrichment analyses

Predicted conserved MBS of rat conserved miRNA families (Targetscan 7.2)^55^, were counted in exclusive gene sets as described in Fig. 2c) and tested for an enrichment against the specific background using a hypergeometric test on the number of binding sites (soon to be made available on Bioconductor).

We considered only those miRNAs, which are considerably expressed in hippocampal neurons at this developmental stage (expressed higher than the median of all detected miRNAs in^15^).

#### 3’UTR length analysis

In order to get an estimate of the average 3’UTR lengths of genes present in somata and processes, genomic coordinates of the longest transcript associated with each detected gene were downloaded in R (AnnotationHub, Ensemble v.100, *Rattus Norvegicus*). As long 3’UTR isoforms are specifically prevalent in the brain^56^, we chose to focus on these in our analysis.

Only 3’UTRs with a length of more than 20 nucleotides were considered for the comparisons.

#### RNA-binding-protein motif analysis

To identify enriched RBP motifs within 3’UTRs in differentially expressed genes, we extracted the matching FASTA sequences according to the criteria specified in the 3’UTR length analysis section. Sequences corresponding to differentially up-/downregulated genes in each compartment upon PTX treatment were submitted to the AME-tool of the meme-suite^57^ in combination with the CISBP database of known RBP motifs^28^.

In AME, standard parameters were used with the *Average Odds Score* as scoring method and *Fisher’s exact test* for the enrichment estimation. RBP-names in the plots were obtained by comparing the AME output motif_ID to the CISBP database motif_ID annotations.

### Graphical representations

Plots were generated using Affinity Designer, Cytoscape & R (ggplot2 (v.3.3.1), ggpubr (v.0.4.0), ggsci (v.2.9), ggrepel (v.0.8.1)).

### Statistics

The number of independent experiments is indicated in the plots, if not explicitly specified differently in the methods section. Boxplots represent median (box: two quantiles around the median; whiskers: Minimal and maximal values; point outside: Outliers). Normality was tested using the Shapiro-Wilk test considering a p-value under 0.05 non-normally distributed. Normally distributed data was tested using one- or two sample Student’s T-test (always two-sided) or ANOVA and otherwise for non-normal data the non-parametric counterpart tests Mann-Whitney-U test or Kruskal-Wallis test. Differences in cumulative distributions was assessed by two-sample Kolmogorow-Smirnow test. Bivariate correlation analysis between two variables was evaluated using the non-parametric Spearman rank correlation using the function ggscatter (from ggpubr).

Additional information about statistical tests not described in the figure legends:

**Figure 3A:**
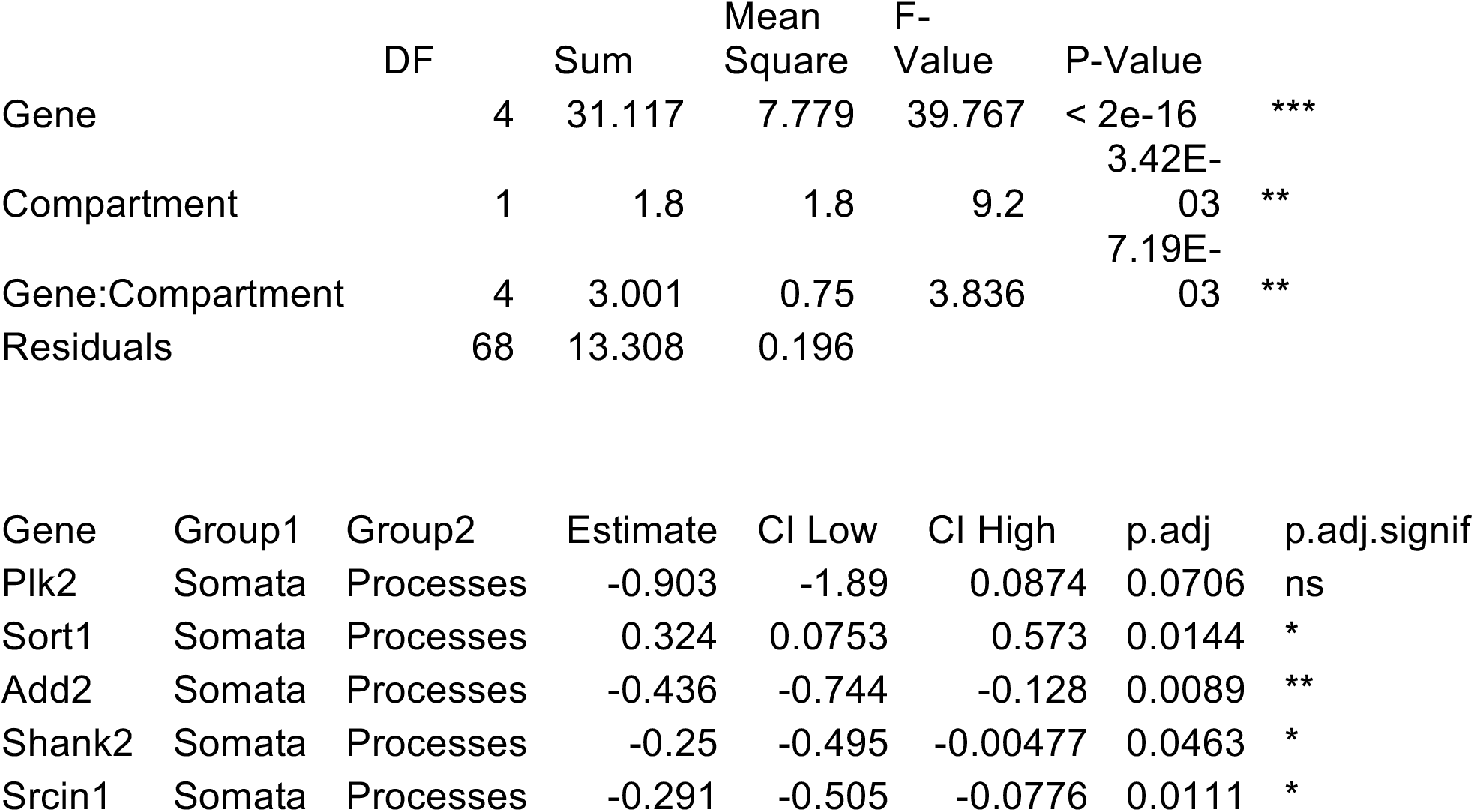
Two-way ANOVA (∼ *Compartment* * *Gene*) with Tukey’s post-hoc test

**Figure 3A:**
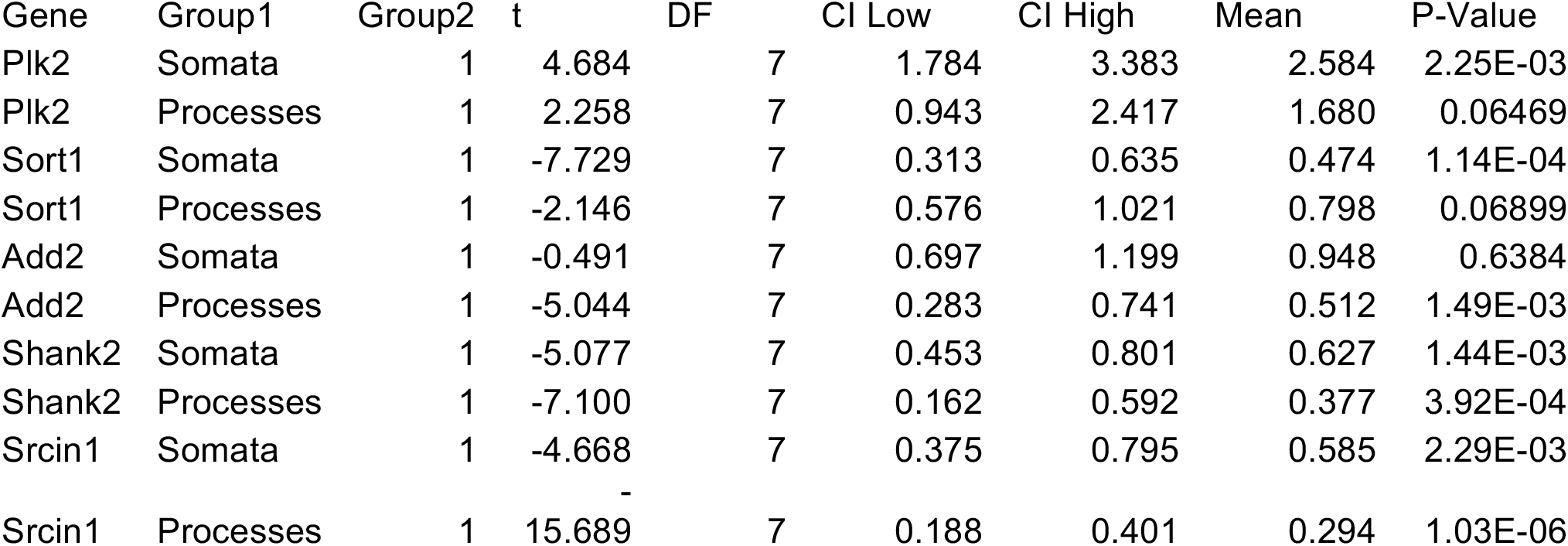
One-sample Student’s T-test (μ_0_=1):

**Figure 3B:**
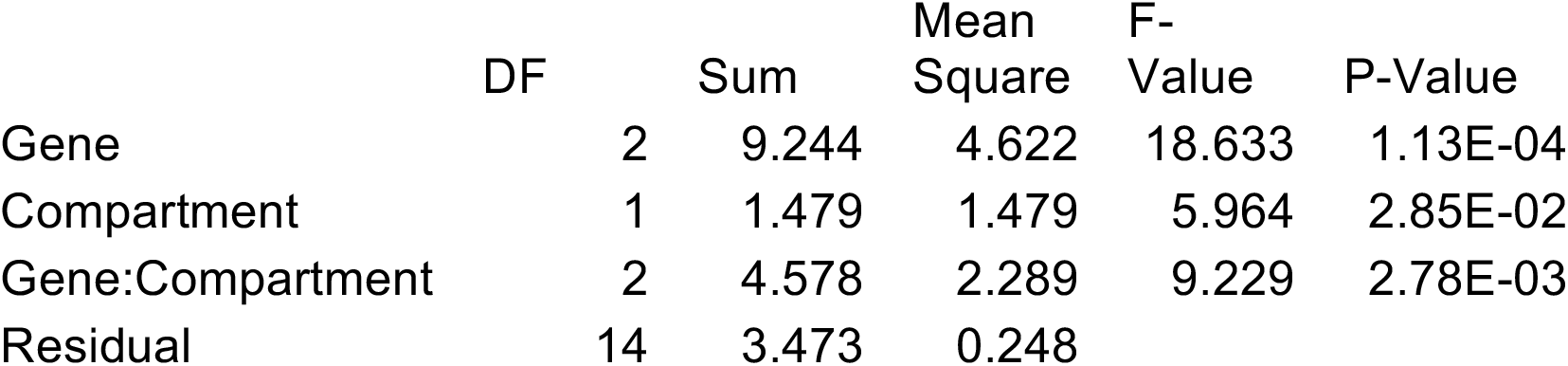
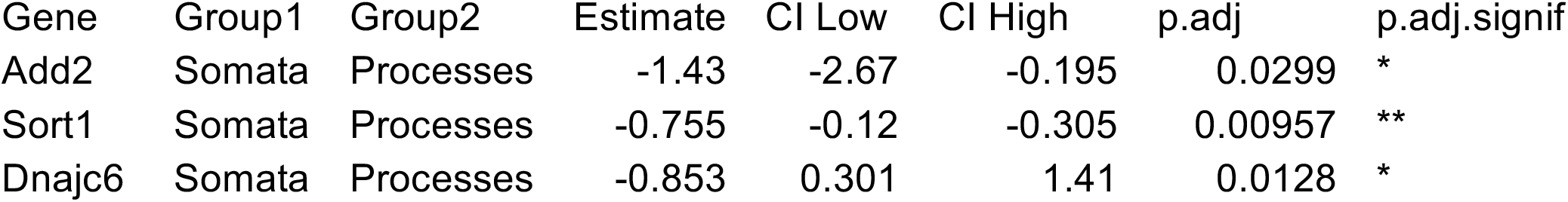
Two-way ANOVA (∼ Compartment * Gene) with Tukey’s post-hoc test

**Figure 5B:**
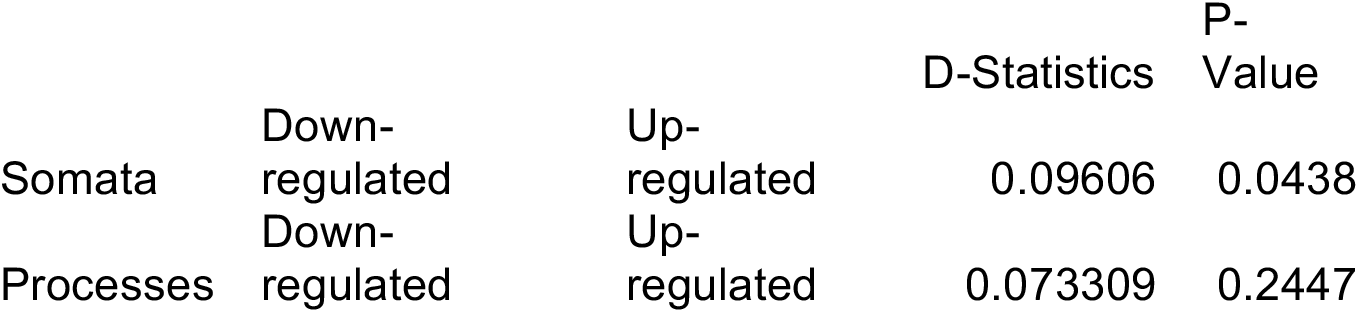
Two-sample Kolmogorov-Smirnov Test (Down-regulated vs Up-regulated in both compartments)

**Figure 5E:**
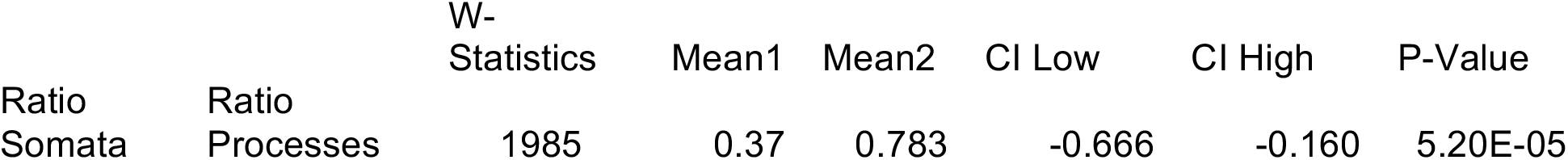
Mann-Whitney U-Test (Ratio in Somata vs Ratio in Processes)

### Real time PCR primers

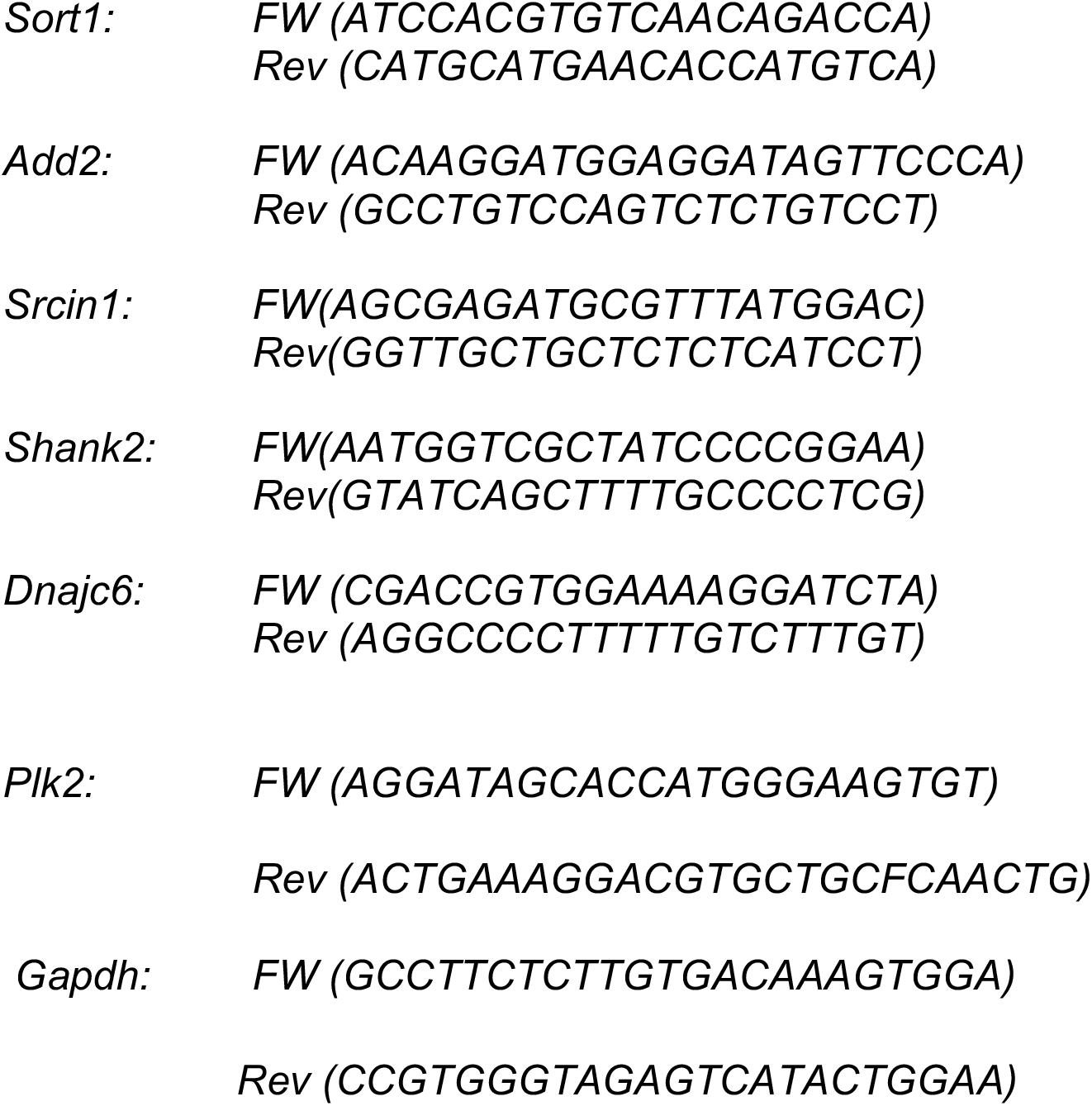

## Acknowledgments

We greatly acknowledge the excellent technical assistance provided by Eva Becker and Tatjana Wüst. This work is funded by a grant from the DFG to CD and GS (SCHR 1136/4-2). The lab of JB is funded by ETH Project Grant ETH-20 19-1, SNSF Grant 310030_172889/1 and the Botnar Foundation.

## Author contributions

DC performed proteomics, qPCR, RNA-seq analysis and generated the figures. MR performed RNA-seq. MS performed motif enrichment analysis. SB and MR performed and analysed the smFISH data. LvZ and JB supervised the proteomics experiments. CD helped with RNA-seq data analysis. PLG supervised RNA-seq and motif enrichment analysis. GS coordinated the project and wrote the manuscript.

## Data availability

RNA-seq data has been deposited to GEO. The mass spectrometry proteomics data have been deposited to the ProteomeXchange Consortium via the PRIDE partner repository. Reviewers will have access to the data via provided passwords/tokens. The following figures have associated raw data: Fig. 1, 2, 4, S1-S7.

