## Supplementary Figures and Legends for "Pervasive compartment-specific regulation of gene expression during homeostatic synaptic scaling"

**Supplementary Figure legends**

**Suppl. Figure 1:** GO-Term analysis of compartment-specific localized genes. Bar graphs illustrating the Top 10 gene ontology terms (MF = molecular function, BP = biological process) of genes enriched in the somata or processes compartment, respectively.

**Suppl. Figure 2:** Principal component analysis of the transcriptomics (A) and proteomics (B) experiments.

**Suppl. Figure 3:** lincRNA and miRNA associated genes are differentially expressed in both compartments upon PTX-treatment. Volcano Plots indicating changes in lincRNA and miRNA genes (Ensembl biotype classification) upon PTX treatment in the somata and processes compartment.

**Suppl. Figure 4:** GO-Term analysis of genes changing upon PTX stimulation in the individual compartments. Depicted are the Top 5 terms of each gene ontology (sorted by p-value, fisher-elim algorithm). Colored bar graphs indicate the number of significantly changing genes associated with each term in the respective compartment (yellow = sign. upregulated genes, red = sign. downregulated genes).

**Suppl. Figure 5:** Volcano plots representing protein down- (red) or up-regulation (yellow) after 48 hours PTX in the somatic and process compartment. FDR < 0.1.

**Suppl. Figure 6**: GO-Term analysis of significantly changing proteins upon PTX stimulation in individual compartments. Shown are the Top 10 significant GO-Terms of the cellular component ontology (sorted by p-value, fisher-elim algorithm) in both compartments. Colored bar graphs indicate the number of significantly changing proteins associated with each term in the respective compartment (yellow = sign. upregulated proteins, red = sign. downregulated proteins).

**Suppl. Figure 7:** miRNA binding site enrichment analysis of all significantly downregulated genes in the processes compartment.

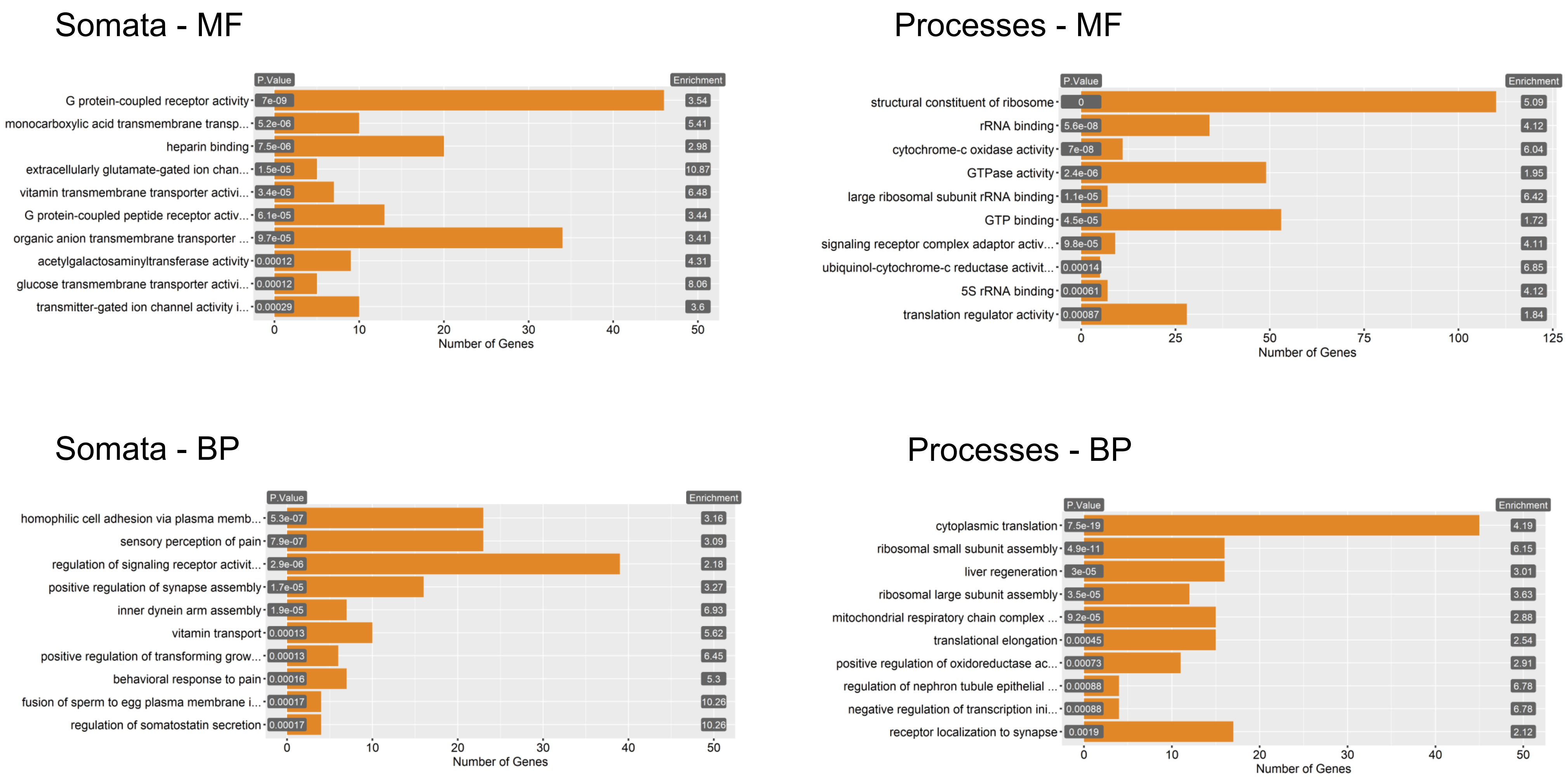
Somata - MF
P.Value
Enrichment
G protein-coupled receptor activity 7e-09 3.54
monocarboxylic acid transmembrane transp... 5.2e-06 5.41
heparin binding 7.5e-06 2.98
extracellularly glutamate-gated ion chan... 1.5e-05 10.87
vitamin transmembrane transporter activi... 3.4e-05 6.48
G protein-coupled peptide receptor activ... 6.1e-05 3.44
organic anion transmembrane transporter ... 9.7e-05 3.41
acetylgalactosaminyltransferase activity 0.00012 4.31
glucose transmembrane transporter activi... 0.00012 8.06
transmitter-gated ion channel activity i... 0.00029 3.6
0 10 20 30 40 50
Number of Genes
Processes - MF
P.Value
Enrichment
structural constituent of ribosome 0 5.09
rRNA binding 5.6e-08 4.12
cytochrome-c oxidase activity 7e-08 6.04
GTPase activity 2.4e-06 1.95
large ribosomal subunit rRNA binding 1.1e-05 6.42
GTP binding 4.5e-05 1.72
signaling receptor complex adaptor activ... 9.8e-05 4.11
ubiquinol-cytochrome-c reductase activit... 0.00014 6.85
5S rRNA binding 0.00061 4.12
translation regulator activity 0.00087 1.84
0 25 50 75 100 125
Number of Genes
Somata - BP
P.Value
Enrichment
homophilic cell adhesion via plasma memb... 5.3e-07 3.16
sensory perception of pain 7.9e-07 3.09
regulation of signaling receptor activit... 2.9e-06 2.18
positive regulation of synapse assembly 1.7e-05 3.27
inner dynein arm assembly 1.9e-05 6.93
vitamin transport 0.00013 5.62
positive regulation of transforming grow... 0.00013 6.45
behavioral response to pain 0.00016 5.3
fusion of sperm to egg plasma membrane i... 0.00017 10.26
regulation of somatostatin secretion 0.00017 10.26
0 10 20 30 40 50
Number of Genes
Processes - BP
P.Value
Enrichment
cytoplasmic translation 7.5e-19 4.19
ribosomal small subunit assembly 4.9e-11 6.15
liver regeneration 3e-05 3.01
ribosomal large subunit assembly 3.5e-05 3.63
mitochondrial respiratory chain complex ... 9.2e-05 2.88
translational elongation 0.00045 2.54
positive regulation of oxidoreductase ac... 0.00073 2.91
regulation of nephron tubule epithelial ... 0.00088 6.78
negative regulation of transcription ini... 0.00088 6.78
receptor localization to synapse 0.0019 2.12
0 10 20 30 40 50
Number of Genes

**Suppl. Fig. S1**

A

Transcriptomics PC1 and PC2

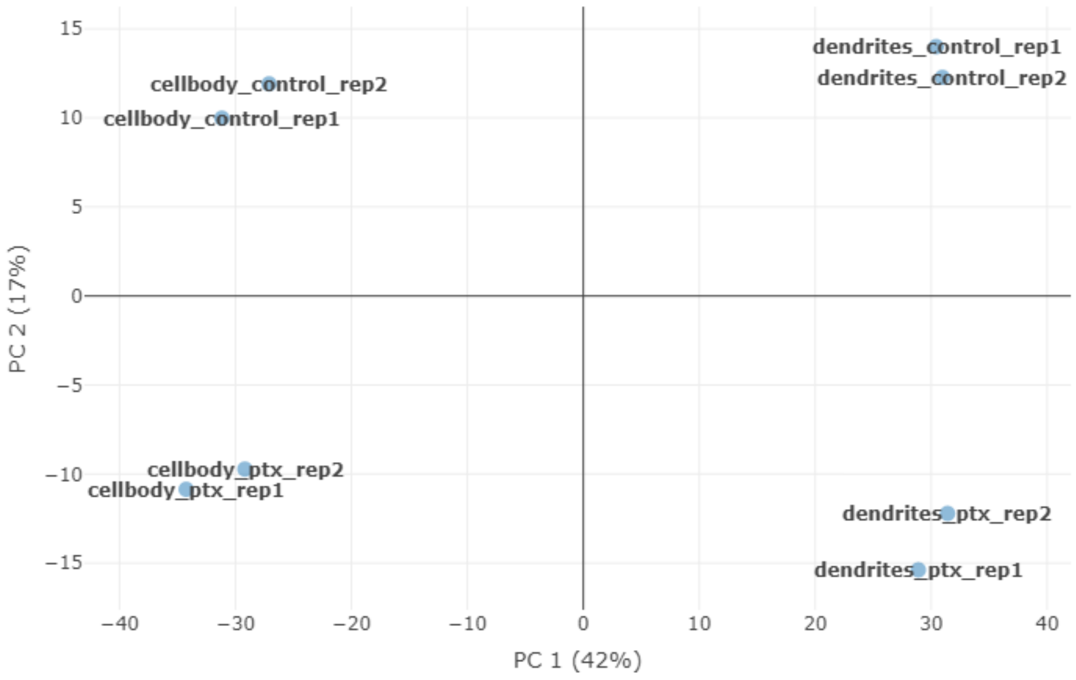


B

Proteomics PC1 and PC2

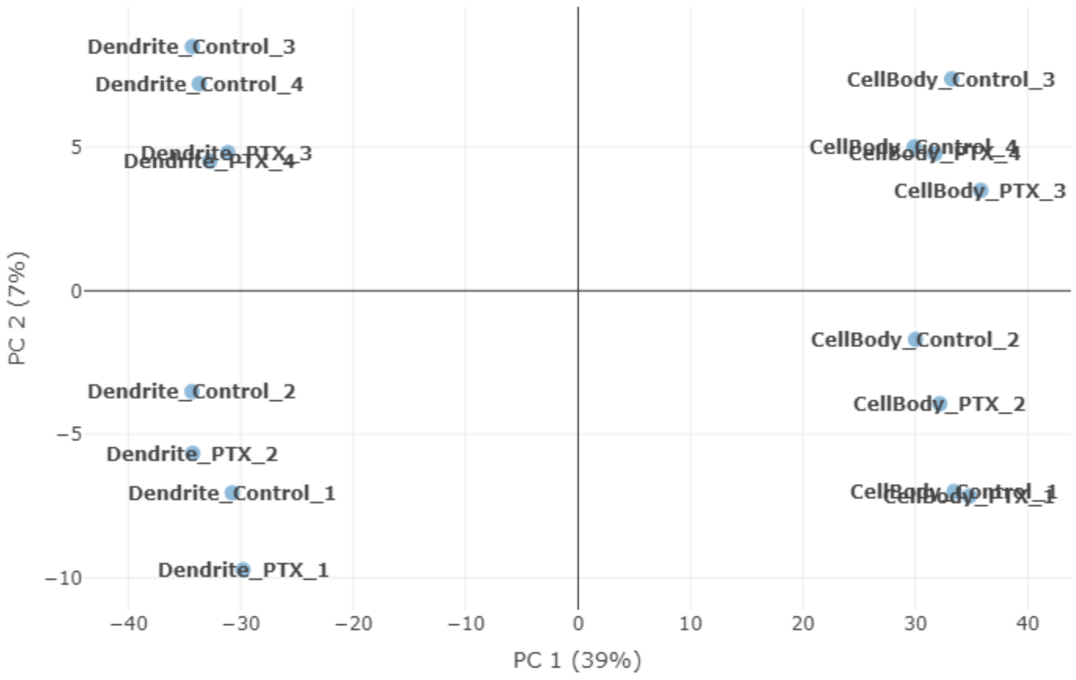


**Suppl. Fig. S2**

### lincRNAs

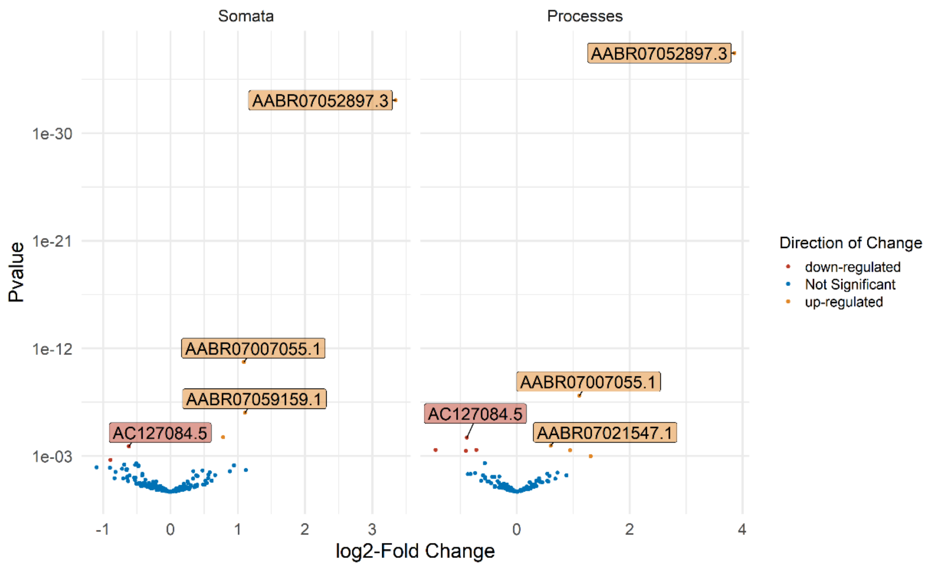


### miRNA genes

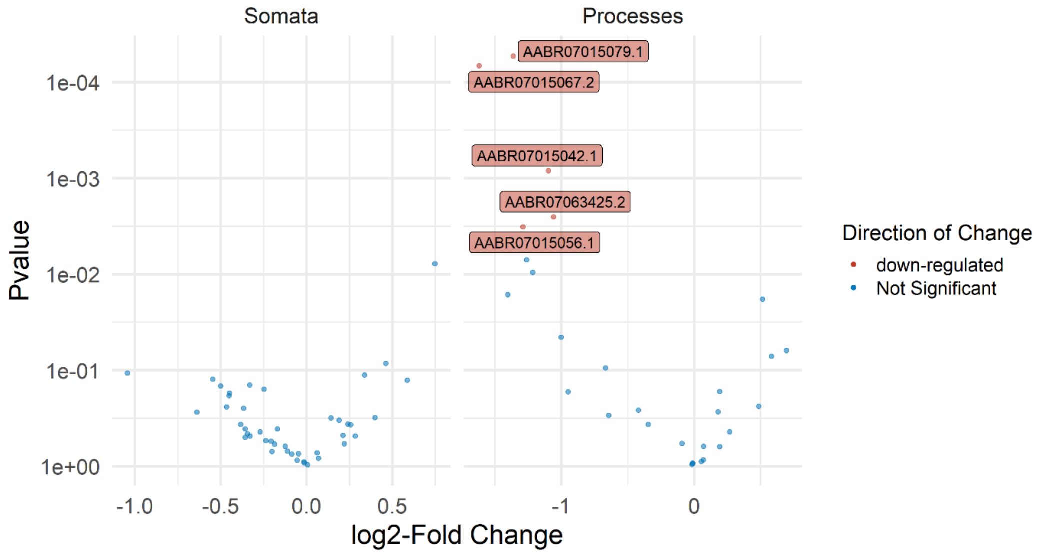


**Suppl. Fig. S3**

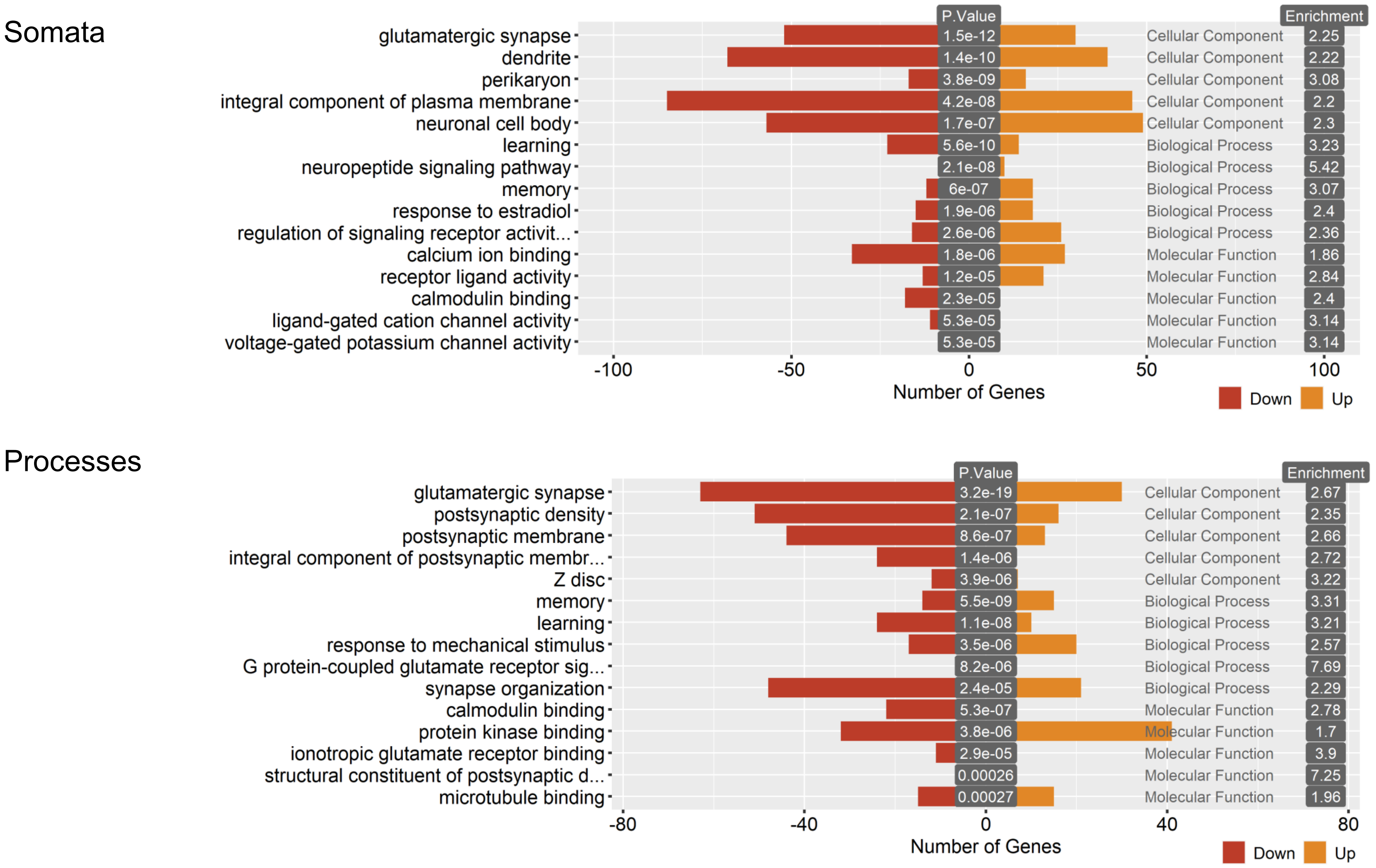
Somata
P.Value
Enrichment
glutamatergic synapse
1.5e-12
Cellular Component
2.25
dendrite
1.4e-10
Cellular Component
2.22
perikaryon
3.8e-09
Cellular Component
3.08
integral component of plasma membrane
4.2e-08
Cellular Component
2.2
neuronal cell body
1.7e-07
Cellular Component
2.3
learning
5.6e-10
Biological Process
3.23
neuropeptide signaling pathway
2.1e-08
Biological Process
5.42
memory
6e-07
Biological Process
3.07
response to estradiol
1.9e-06
Biological Process
2.4
regulation of signaling receptor activit...
2.6e-06
Biological Process
2.36
calcium ion binding
1.8e-06
Molecular Function
1.86
receptor ligand activity
1.2e-05
Molecular Function
2.84
calmodulin binding
2.3e-05
Molecular Function
2.4
ligand-gated cation channel activity
5.3e-05
Molecular Function
3.14
voltage-gated potassium channel activity
5.3e-05
Molecular Function
3.14
-100
-50
0
50
100
Number of Genes
Down
Up
Processes
P.Value
Enrichment
glutamatergic synapse
3.2e-19
Cellular Component
2.67
postsynaptic density
2.1e-07
Cellular Component
2.35
postsynaptic membrane
8.6e-07
Cellular Component
2.66
integral component of postsynaptic membr...
1.4e-06
Cellular Component
2.72
Z disc
3.9e-06
Cellular Component
3.22
memory
5.5e-09
Biological Process
3.31
learning
1.1e-08
Biological Process
3.21
response to mechanical stimulus
3.5e-06
Biological Process
2.57
G protein-coupled glutamate receptor sig...
8.2e-06
Biological Process
7.69
synapse organization
2.4e-05
Biological Process
2.29
calmodulin binding
5.3e-07
Molecular Function
2.78
protein kinase binding
3.8e-06
Molecular Function
1.7
ionotropic glutamate receptor binding
2.9e-05
Molecular Function
3.9
structural constituent of postsynaptic d...
0.00026
Molecular Function
7.25
microtubule binding
0.00027
Molecular Function
1.96
-80
-40
0
40
80
Number of Genes
Down
Up

**Suppl. Fig. S4**

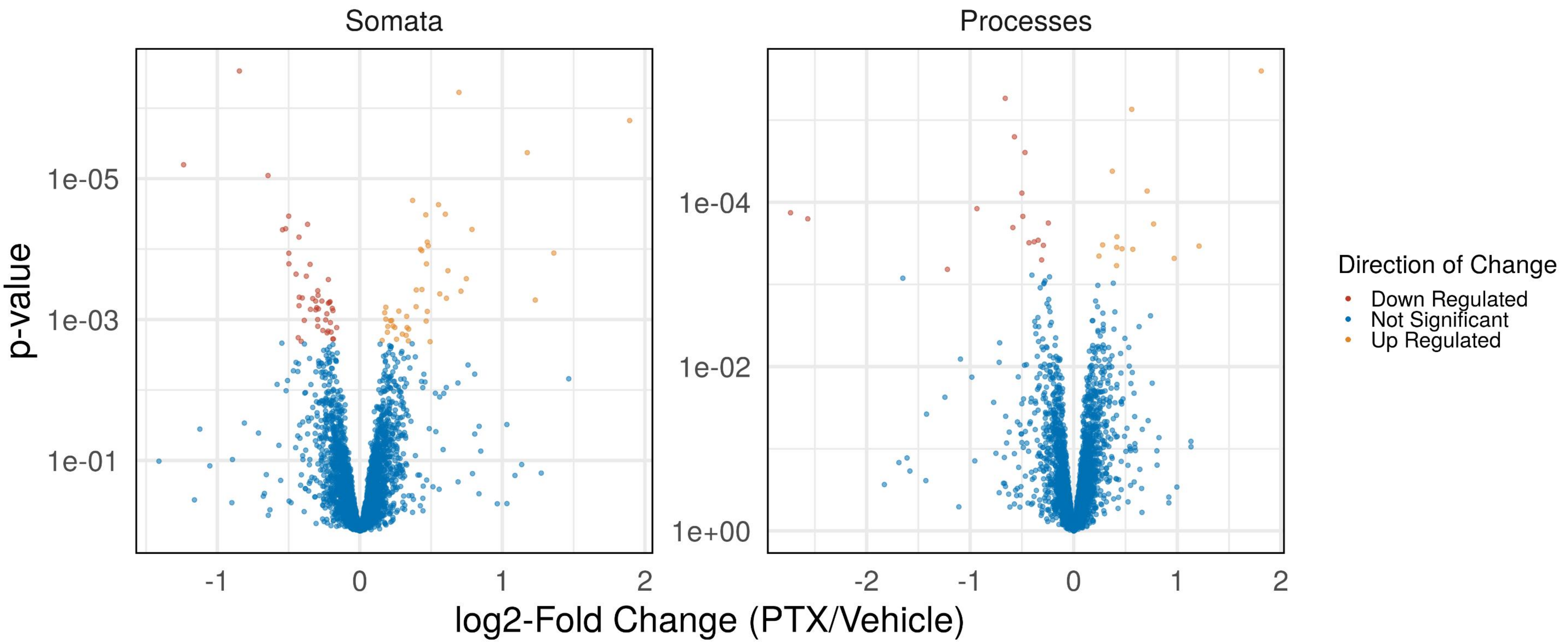
Somata
Processes
p-value
1e-05
1e-03
1e-01
1e-04
1e-02
1e+00
-1
0
1
2
-2
log2-Fold Change (PTX/Vehicle)
Direction of Change
Down Regulated
Not Significant
Up Regulated

**Suppl. Fig. S5**

Somata

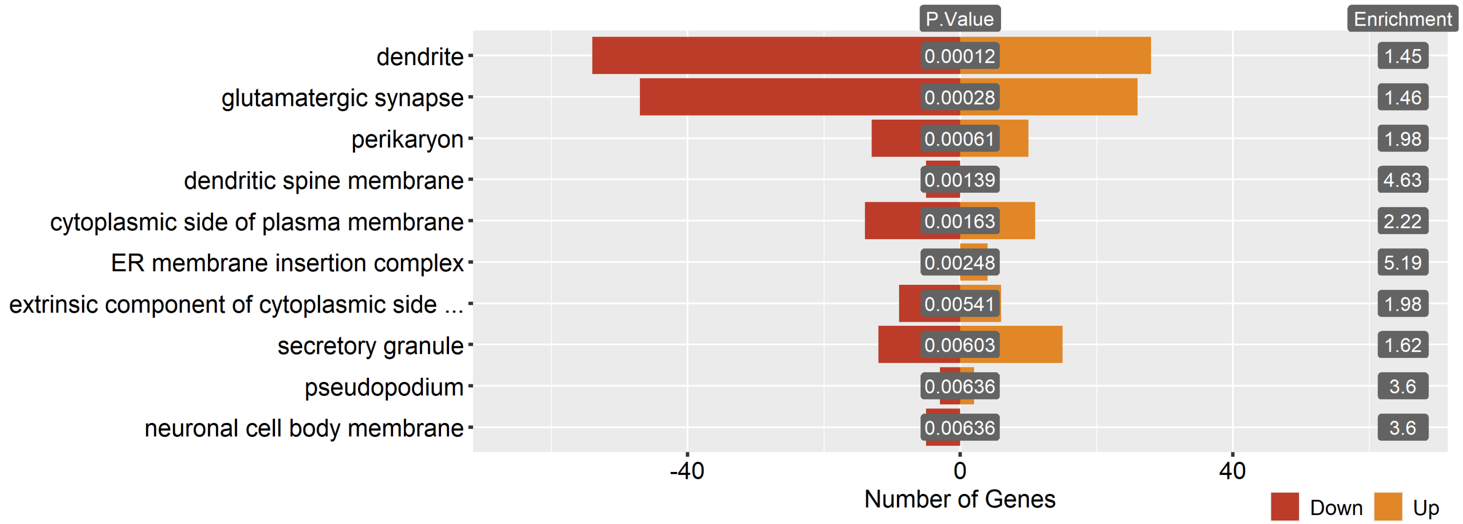


Processes

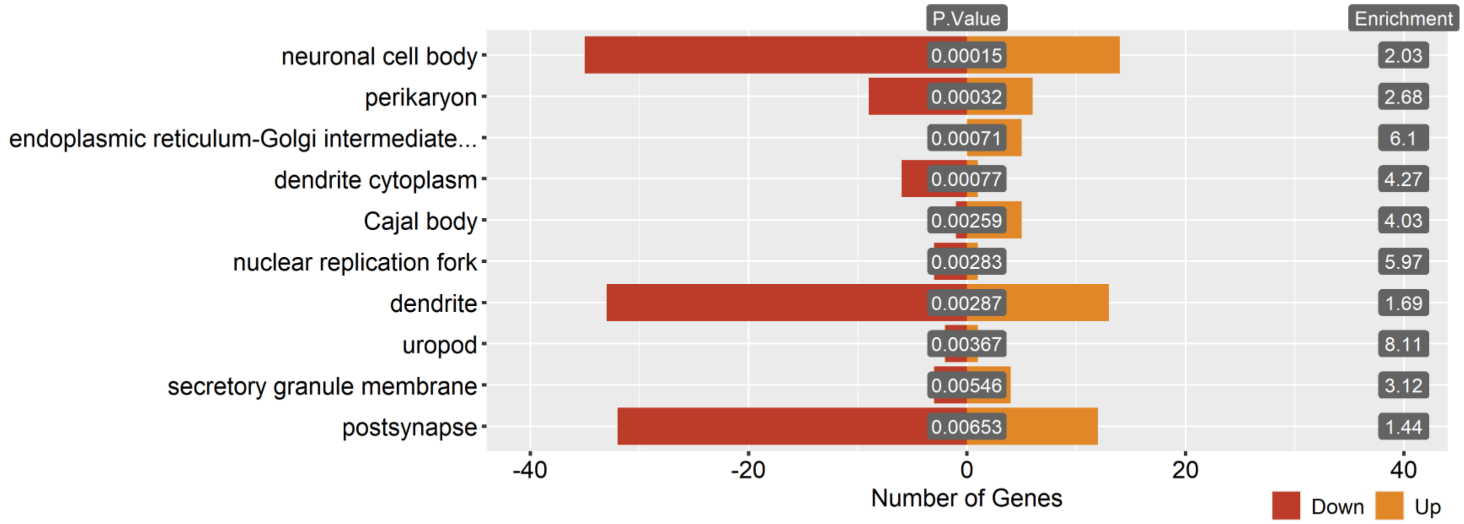


**Suppl. Fig. S6**

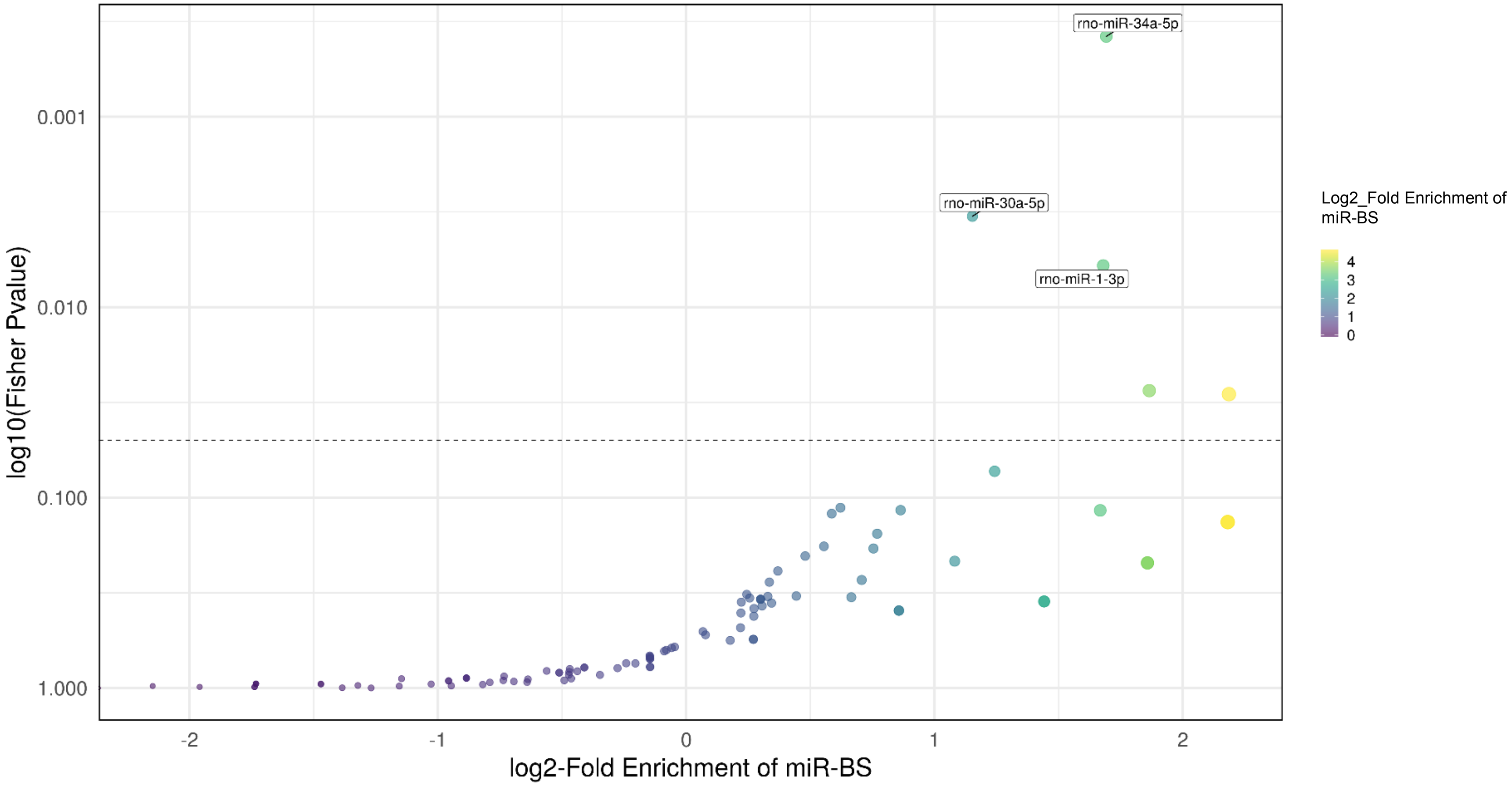
rno-miR-34a-5p
rno-miR-30a-5p
rno-miR-1-3p
0.001
0.010
0.100
1.000
log10(Fisher Pvalue)
-2
-1
0
1
2
log2-Fold Enrichment of miR-BS
Log2_Fold Enrichment of miR-BS
4
3
2
1
0

**Suppl. Fig. S7**
